# The impact of light and thioredoxins on the plant thiol-disulfide proteome

**DOI:** 10.1101/2023.10.17.562744

**Authors:** Liang-Yu Hou, Frederik Sommer, Louis Poeker, Dejan Dziubek, Michael Schroda, Peter Geigenberger

## Abstract

Thiol-based redox regulation is a crucial post-translational mechanism to acclimate plants to changing light availability. Here, we conduct a biotin-switch-based redox proteomics study to systematically investigate dynamics of the thiol-redox network in response to temporal changes in light availability and across genotypes lacking parts of the thioredoxin (Trx) or NADPH-Trx-reductase C (NTRC) systems in the chloroplast. Time-resolved dynamics revealed light leading to marked decreases in the oxidation states of many chloroplast proteins with photosynthetic functions during the first 10 min, followed by their partial re-oxidation after 2-6 hours into the photoperiod. This involved *f*, *m* and *x*-type Trx proteins showing similar light-induced reduction-oxidation dynamics, while NTRC, 2-Cys-Prx and Trx *y*2 showed an opposing pattern, being more oxidized in the light than the dark. In Arabidopsis *trxf1f2*, *trxm1m2* or *ntrc* mutants, in the light most proteins showed increased oxidation states than wild type, suggesting their light-dependent dynamics being related to the NTRC/Trx networks. While NTRC deficiency had a strong influence in all light conditions, deficiencies in *f*- or *m*-type Trxs showed differential impacts on the thiol-redox proteome depending on the light environment, being higher in constant or fluctuating light, respectively. Results indicate plant redox proteomes to be subject to dynamic changes in reductive and oxidative pathways to cooperatively fine-tune photosynthetic and metabolic processes in the light. This involves *f*-type Trxs and NTRC to play a role in constant medium light, while both *m*-type Trxs and NTRC being important to balance changes in protein redox-pattern during dynamic alterations in fluctuating light intensities.

**One sentence summary:** The plant protein redoxome shows light-dependent reduction and reoxidation dynamics linked to Trxs *f*1/*f*2, *m*1/*m*2 and NTRC, being of different importance depending on the extent of light variability.

## Introduction

Due to their sessile lifestyle, plants experience countless environmental challenges. Among these environmental stimuli, light availability is probably the most crucial factor, since plants rely on photosynthesis to sustain their development and growth. In nature, the fluctuations of light availability occur on a time-scale of seconds to months and encompass large changes in irradiance. As sessile organisms, plants have evolved sophisticated strategies to rapidly acclimate to frequently changed light intensities. While proteomics studies show that there are only very minor changes in the abundance of proteins in response to fluctuating light (FL; Niedermaier et al., 2020) or during diurnal changes in light availability(Uhrig et al., 2021), the core of these regulatory mechanisms to rapidly acclimate light fluctuations is thiol-based redox regulation of proteins. This kind of redox regulation exclusively occurs at thiol groups of cysteine residues, which further changes protein activity and conformation (Cremers and Jakob, 2013). The reactions of such disulfide-dithiol interchange rely on a group of proteins called thioredoxins (Trxs; Meyer et al., 2012).

Thioredoxins are 12-14 kDa proteins harboring a conserved active site, WCGPC (Holmgren, 1995). Trxs usually serve as disulfide reductase to reduce and activate their target enzymes using NADPH via the mediation of NADPH-dependent Trx reductase (NTR) proteins (Jacquot et al., 2009). In heterotrophic organisms, there are usually one or two genes encoding Trx and NTR proteins in governing numerous redox regulations (Meyer et al., 2012). Intriguingly, autotrophic organisms have rather complicated redox networks. For example, cyanobacteria, algae and plants usually constitute a great number of Trx and NTR proteins (Geigenberger et al., 2017; Zaffagnini et al., 2019). Unlike Trx proteins of heterotrophic organisms exclusively gaining reducing power from NTR proteins, the chloroplastic Trxs of autotrophic organisms can also drain reducing equivalents from photosynthetically reduced ferredoxin (Fdx) with the involvement of Fdx-dependent Trx reductase (FTR) proteins (Schürmann and Buchanan, 2008). In Arabidopsis, there are twenty Trxs distributed in different subcellular compartments (Geigenberger et al., 2017; Zaffagnini et al., 2019). Among these Trx proteins, the chloroplastic Trxs are of great importance for chloroplast metabolism and light acclimation. With analyses in Trx mutant lines, the *f*-type Trxs were determined as positive regulators for light-dependent carbon fixation, specifically during dark-light transitions (Michelet et al., 2013; Thormählen et al., 2013; Thormählen et al., 2015; Naranjo et al., 2016a). The *m*-type Trxs were found to regulate the redox state of NADP-dependent malate dehydrogenase (NADP-MDH) to modulate the shuttle of reducing equivalents via the malate valve (Okegawa and Motohashi, 2015; Thormählen et al., 2017; Selinski and Scheibe, 2019). The *x*-type and *y*-type Trxs were found to be involved in antioxidation processes (Collin et al., 2004; Lamkemeyer et al., 2006; Navrot et al., 2006; Bohrer et al., 2012), while the *z*-type Trx was proposed to regulate plastidial gene expression (Arsova et al., 2010).

In addition to the Fdx-Trx system, which is directly linked to light, a unique NTR protein (NTRC) with joint Trx domain at its C-terminal end was discovered to use NADPH to reduce the hydrogen peroxide scavenging enzyme 2-Cys peroxiredoxin (2-Cys Prx) in the chloroplast (Serrato et al., 2004). Interestingly, NTRC was also found to affect broad chloroplast metabolism, including the redox states of plastidial Trx targets (Serrato et al., 2004; Michalska et al., 2009; Lepistö et al., 2013; Richter et al., 2013; Pérez-Ruiz et al., 2014; Thormählen et al., 2015; Carrillo et al., 2016; Naranjo et al., 2016b; Da et al., 2017). These effects were shown to be mainly indirect, with the redox balance of 2-Cys Prx indirectly modulating Trx-regulated enzymes (Pérez-Ruiz et al., 2017; Cejudo et al., 2019; Cejudo et al., 2021; Lampl et al., 2022). However, a comprehensive analysis of the redox proteome to investigate the roles of NTRC and Fdx-Trxs in more detail remains to be studied.

To further resolve the Trx-mediated redox network, identifying the downstream targets of Trxs became a much-anticipated research topic. Over the past two decades, researchers have made a great effort on identifying Trx-target proteins. The initial approach to pinpoint Trx targets relies on the basis of disulfide-dithiol interchange taking place between Trx and its target protein. It has been established that the N-terminal cysteine of Trx active site first reacts with the disulfide bond of the target protein, leading to the formation of a transient heterodimer. The C-terminal cysteine of Trx active site subsequently initiates a second reaction at the target disulfide bond to resolve this heterodimer. Afterward, the oxidized Trx and the reduced target protein are further dissociated (Brandes et al., 1993; Holmgren, 1995). Therefore, substitution of the C-terminal active cysteine to another amino acid will disrupt the dissociation between Trx and its target, stabilizing the heterodimer. This monocysteinic Trx variant can serve as bait to pull down its interacting targets. Such technique together with two-dimensional gel electrophoresis has been extensively applied to isolate Trx targets in cyanobacteria (Lindahl and Florencio, 2003; Pérez-Pérez et al., 2006), Chlamydomonas (Goyer et al., 2002; Lemaire et al., 2004) and many land plants (Balmer et al., 2003; Balmer et al., 2004b; Balmer et al., 2004a; Wong et al., 2004; Yamazaki et al., 2004; Marchand et al., 2006; Bartsch et al., 2008; Montrichard et al., 2009; Marchand et al., 2010; Yoshida et al., 2013) in vitro. Indeed, these studies identified hundreds of potential Trx target proteins that require further confirmation and evaluation, specifically to demonstrate their importance in vivo or to exclude unspecific binding due to Trxs acting as chaperones.

Another commonly used approach to identify Trx targets is to label the thiol group of cysteine using a redox-active probe (Yano et al., 2001; Balmer et al., 2006; Alkhalfioui et al., 2007; Hall et al., 2010). In addition to the qualitative application, several quantitative modifications of this strategy such as biotinylated iodoacetamide (BIAM) switch assay and an integrative proteomics method using cysteine-reactive isobaric tandem mass tags (CysTMT, iTRAQ) differential cysteine labeling in combination with gel shifts were also implemented to evaluate changes in the thiol redox status of proteins in various conditions (Parker et al., 2015; Pérez-Pérez et al., 2017; Löwe et al., 2019; Zimmer et al., 2021).

Although a wide variety of redox proteomics approaches have been well developed, a quantitative analysis of the thiol-disulfide redox proteome in response to different light intensities is somewhat scarce. In the current study, we implemented the biotin switch assay together with label-free quantification to evaluate the relative changes of protein oxidation levels at different time points into the photoperiod and during fluctuating light intensities in Arabidopsis plants. To further investigate the impact of the chloroplast thiol-redox network on changes in protein oxidation levels during light acclimation, we analyzed several well-characterized Trx mutants including *trxf1f2* (Naranjo et al., 2016a) and *trxm1m2* double mutants (Thormählen et al., 2017) as well as the *ntrc* single mutant (Serrato et al., 2004) in comparison to the wild type. By investigating light-dependent dynamics in the protein redoxome, we demonstrated that large sets of proteins involved in photosynthetic light reactions, Calvin Benson Cycle (CBC) and carbohydrate metabolism are reduced within 10 minutes of illumination, while they are subject to re-oxidation processes after 2-6 h into the light period. Interestingly, *f*, *m* and *x*-type Trx proteins showed similar light-induced reduction-oxidation dynamics as their photosynthetic targets, while NTRC, 2-Cys-Prx and Trx *y*2 showed an opposing pattern, being more oxidized in the light, compared to the dark. Through studying the protein redoxome in the mutant lines deficient in parts of the thiol-disulfide system, we uncovered that light-dependent *f*- and *m*-type Trxs play distinct roles in modulating protein oxidation states in different light conditions. While Trxs *f*1/*f*2 are more important during regular growth light and in the high-light phases of fluctuating light (FL), Trxs *m*1/*m*2 mainly play a role during the low-light FL phases. In contrast, NTRC was found to be indispensable to modulate the oxidation-state of photosynthetic proteins in all light conditions, probably due to its role to regulate oxidative signals depending on NADPH. In addition to well-known photosynthetic targets, we also identified proteins involved in antioxidation processes and metabolism of amino acids and proteins to be subject to light-and Trx-dependent redox modulation.

## Results

### Light leads to temporal and spatial dynamics of the leaf redox proteome involving mainly proteins with photosynthetic functions located in the plastid

To understand effects of illumination on leaf protein redox states, we performed a time-resolved study in the wild type. The design of this experiment is outlined in Supplemental Figure S1A. Arabidopsis plants were grown under medium light intensity (ML, 150 μmol photons m^-2^ s^-1^ with a 12-h-dark/12-h-light regime; 22°C) for four weeks, before leaf samples were harvested at different diurnal time points by freezing whole rosettes directly into liquid nitrogen. We first harvested leaf samples at the end of the night (EN, dark conditions), which served as the control. The following time points were 10 min (L10), 120 min (L120) and 360 min (L360) into the photoperiod, representing the effects of short, mid and long-term illumination (Supplemental Fig. S1A). To analyze proteins that show light-dependent changes in their oxidation states, leaf samples were subsequently subjected to a redox-proteomics method, which is described in Supplemental Figure S1B. After protein extraction in the presence of N-ethylmaleimide (NEM) to alkylate (and block) the free thiol residues of cysteines, oxidized disulfides were subsequently reduced by DTT treatment, and the resulting free thiol residues labeled with a redox-active biotin. The biotinylated proteins were further isolated using a streptavidin resin, while mass spectrometry was used for protein identification and quantification (Supplemental Fig. S1B). It must be noted, as we used NEM to alkylate free thiol residues, our method mainly detected proteins subject to disulfide rather than sulfenic acid modifications.

Using this approach, we successfully identified 1980 proteins which showed light-induced changes in their oxidation states. Subsequently, proteins that have been detected in less than three biological replicates were considered as low-abundance targets, and omitted from the following data processing, as these identified targets were not valid for statistical analyses. Then the remaining 1038 proteins were subjected to statistical analyses in comparison to the EN (dark) samples. To do this, we calculated the fold changes of oxidation levels between illuminated and dark samples to evaluate the effects of illumination on protein oxidation states. Furthermore, we preformed ANOVA with Dunnett’s test, which yields a probability value (P-value) to determine if the changes are statistically significant. Using these criteria, 319 proteins were finally identified to harbor statistically significant (P<0.05) changes in oxidation states in response to illumination and selected for the following analyses (Supplemental Table S1).

Because our approach only detects the oxidized forms of the proteins, but not their reduction levels, calculation of absolute reduction/oxidation ratios was not feasible. We therefore calculated the protein oxidation states at different time points into the photoperiod relative to EN conditions. This approach is subject to possible errors, if there are simultaneous changes in overall protein abundance during the dark-to-light transients. However, a comprehensive proteomics study in Arabidopsis during the diurnal cycle showed that only around 6% of the quantified proteome revealed marked changes in abundance over the course of a day (Uhrig et al., 2021). By comparing the proteins identified in our present study (Supplemental Table S1) with the published data set of Uhrig et al. (2021), we found that only very few of the proteins that were identified to be subject to changes in oxidation levels (16 out of 319) were also reported to be subject to diurnal changes in overall protein levels (Supplemental Table S2). These 16 proteins listed in Table S2 were not in the focus of our study. Overall this shows that, compared to redox alterations, protein abundance changes are of minor importance under these conditions. Our approach is therefore appropriate to evaluate dark-to-light changes in protein redox states during a time course of minutes to hours.

To get a first overview, we grouped the 319 proteins according to their subcellular localization and biological functions (Supplemental Table S1). As shown in Figure 1A, around 39% of identified proteins are localized in the plastid; 21% in the cytosol and 9% in mitochondria, while 31% are distributed to various other subcellular compartments. When looking at their annotated functions, 13% of identified proteins are involved in photosynthesis, 4% in cellular respiration, 10% in various other metabolic processes (e.g., carbohydrate, amino acid, lipid metabolism) and 7% in redox homeostasis. Interestingly, a majority of 19% of identified proteins is involved in RNA and protein processes. The rest (47% of identified proteins) are allocated to the group of other cellular processes and unknown function (Figure 1B).

**Figure 1.**
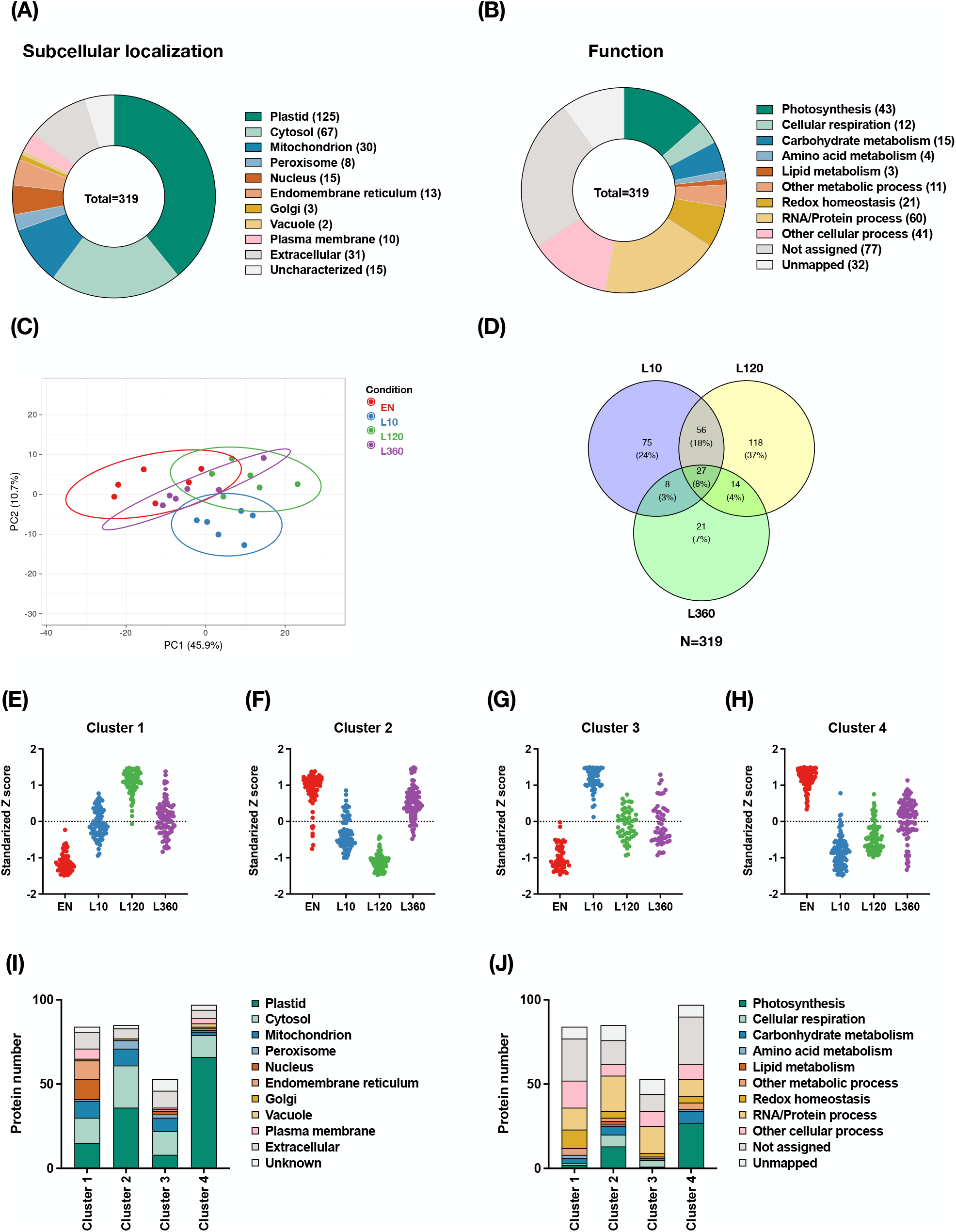
Light-dependent dynamics in the landscape of protein-oxidation changes at different time points into the photoperiod. Redox proteomics analyses of Arabidopsis plants sampled 10 (L10), 120 (L120) and 360 min (L360) into the photoperiod, compared to end of the night (EN, dark conditions). **(A)** Subcellular localization of proteins showing significant changes in their oxidation levels in the light, compared to the dark. **(B)** Functional categories of proteins showing significant changes in their oxidation levels in the light, compared to the dark. **(C)** Principal component analysis displaying the distinctions of protein oxidation states between EN, and L10, L120 and L360. **(D)** Venn diagram highlighting the distribution of proteins subject to significant changes in oxidation states at different time points into the photoperiod, compared to EN. **(E)** - **(H)** Unsupervised cluster analysis showing the grouping of proteins that display different redox changes in response to a time course in the light. **(I)** The subcellular localization of proteins from the different clusters. **(J)** The biological function categories of proteins from the different clusters. Data are based on three to six biological replicates.

To further evaluate light-dependent effects on protein redox states, we performed a principal component analysis (PCA). Results show that the L10 samples were clearly deviated from the EN (dark) samples, while L120 and L360 samples showed a progressive overlap with EN (Fig. 1C). This indicates that protein oxidation states changed dramatically within the first 10 min of illumination, followed by a partial recovery at later time points. The latter increased progressively with time, being more complete after 360, compared to 120 min of illumination (Fig. 1C). The Venn diagram (Fig. 1D) shows the number of proteins with different redox states after different time points of illumination, compared to EN (dark) conditions. The non-overlapping regions of L10, L120 and L360 sample sets comprised 24%, 37% and 7% of the 319 proteins showing differential oxidation states, respectively. The overlapping region between L10 and L120 comprised 18% of total identified targets, indicating that the changes of oxidation state in these proteins endure in the short and medium-term illumination. Rather few targets located in the overlapping region between L360 and L10 (3% of total) as well as L360 and L120 (4% of total) suggesting that the effects of longer-term light exposure on protein redox states are relatively minor when compared to short-term and medium-term illumination (Figure 1D). It is worth noting that there were 27 proteins (8% of total) displaying significant redox changes in all three time points of illumination compared to EN (dark), including Calvin-cycle protein 12 (CP12), NADP-dependent malate dehydrogenase (MDH) and POSTILLUMINATION CHLOROPHYLL FLUORESCENCE INCREASE (PIFI). In these proteins, light-induced changes in the oxidation status were maintained independent of the duration of illumination (Supplemental Table S3).

To further visualize the redox patterns of identified targets, we conducted an unsupervised cluster analysis. The 319 target proteins showing significant changes in redox state were categorized into four clusters. Clusters 1 and 3 contained the targets showing a strong increase in oxidation within the first 10 min (cluster 3) or 2 h of light exposure (cluster 1), while there was a partial re-reduction at later time points (Fig. 1, E and G). These mainly included proteins located outside the plastid (Fig. 1I) and involved in functional categories outside of photosynthesis, such as other metabolic processes, redox-homeostasis and RNA/protein processes (Fig. 1J). Targets in clusters 2 and 4 instead showed an opposite pattern, with proteins getting strongly reduced within 10 min (cluster 4) or 2 h upon illumination (cluster 2), while they underwent some re-oxidation at later time points (Fig. 1, F and H). Notably, most plastidial targets (Fig. 1I) and proteins involved in photosynthetic processes were categorized to clusters 2 and 4 (Fig. 1J), the percentage of these proteins being highest in cluster 4 (Fig. 1, I and J). This implies that a large part of plastidial proteins, especially those participating in photosynthesis, becomes reduced within the first 10 min of light exposure, as a possible “kick off” signal to activate photosynthetic processes. A partial re-oxidation occurs at later time points, specifically after 360 min, indicating that during long-term light exposure oxidative processes come into play.

### Light leads to a rapid increase in the reduction of proteins associated to photosynthesis followed by their partial re-oxidation at later time points

To in-depth understand the effects of light exposure on protein oxidation states in the wild type, we categorized the targets showing significant changes into more detailed functional groups and evaluated their protein oxidation levels (Fig. 2). Within the first 10 min, light led to a rapid decrease in oxidation levels of almost all identified proteins involved in photosynthetic light reactions (Fig. 2A), CBC (Fig. 2B) and major CHO metabolism (Fig. 2C). As shown in Figure 2A, 2B and 2C, these included 20 proteins of light reactions (i.e. subunits of ATP-synthase, PSI and PSII reaction center, Chl a-b binding proteins, NDH, PIFI and protein curvature thylakoid 1 B), 14 proteins associated to key steps of the CBC, and 19 proteins of major CHO metabolism, specifically starch and hexose-phosphate metabolism. Intriguingly, only two target proteins listed within these categories, PsbP domain-containing protein 6 (PPD6) and root isoform of FNR (RFNR1) behaved in an opposing manner, being oxidized upon illumination, instead of being reduced (Fig. 2C), the RFNR1 being not directly involved in photosynthetic metabolism. It is worth noting that RFNR harbors opposite properties compared to LFNR. RFNR uses NADPH derived from the oxidative pentose phosphate pathway (OPPP) to reduce Fdx, which further offers reducing equivalents to the enzymes involved in nitrogen metabolism (Hanke et al., 2005). Thus, the oxidation of RFNR1 might contribute to the active transferring of reducing power to Fdx and downstream enzymes. These results reveal a whole set of photosynthetic proteins related to light reaction, CBC and major CHO metabolism being subject to rapid light-dependent reduction. Indeed, all identified proteins associated to the CBC showed a marked light-dependent decrease in oxidation state within the first 10 min of illumination (Fig. 2B), which most likely involves the Fdx-Trx system (Schürmann and Buchanan, 2008; Yoshida et al., 2022). In confirmation of this, as indicated in Figure 2, all CBC proteins and a large part of the identified proteins associated to light reactions and plastidial CHO metabolism have been reported as Trx targets in previous studies (Lindahl and Kieselbach, 2009).

**Figure 2.**
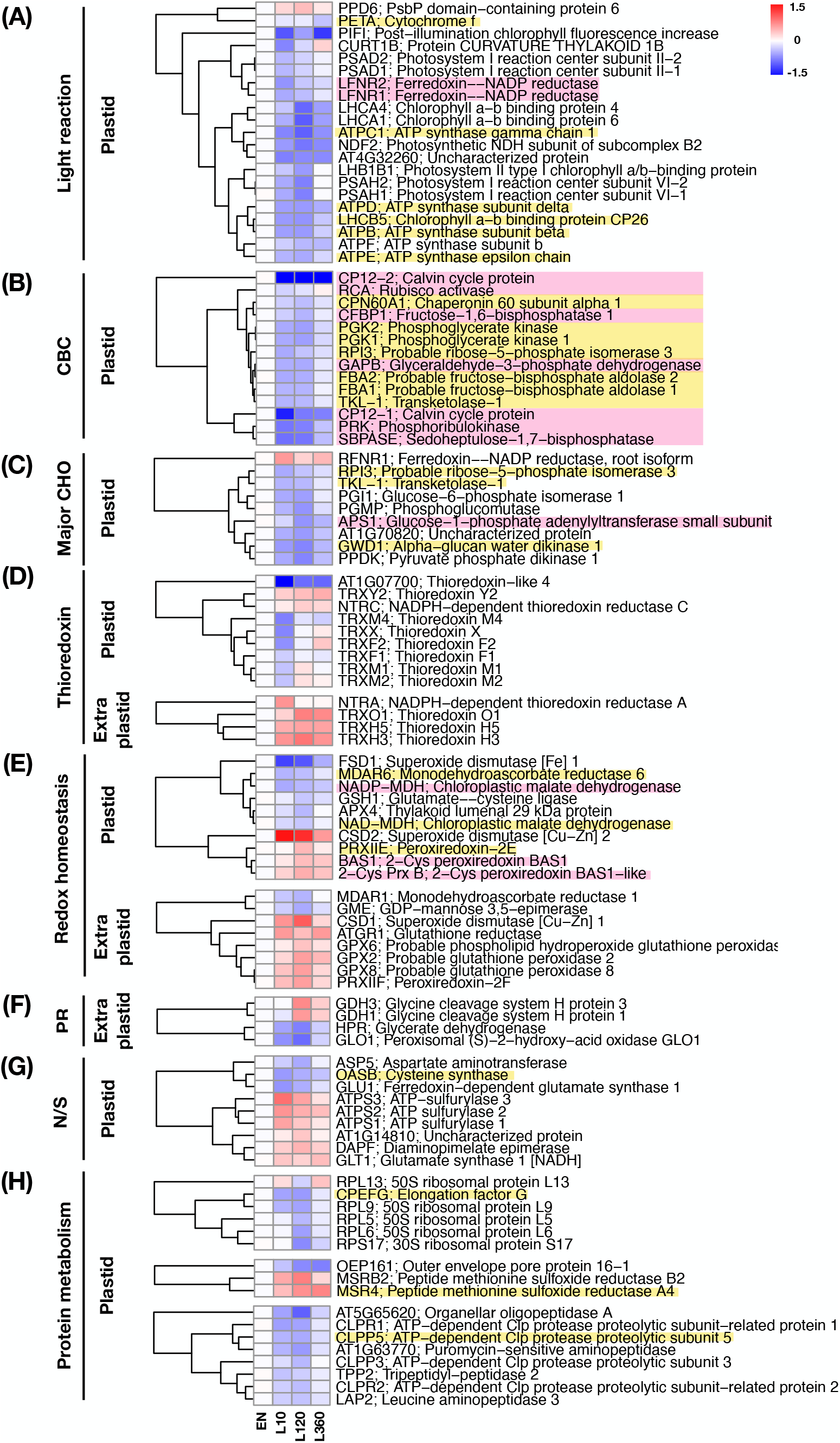
Protein-oxidation changes in photosynthetic processes, redox balance, metabolic pathways and protein metabolism in response to light. The heatmaps summarize the log_2_ fold changes of protein oxidation levels at 10 (L10), 120 (L120) and 360 min (L360) into the photoperiod, relative to the end of the night (EN, dark conditions). Proteins were allocated to different groups of biological functions including **(A)** light reaction, Calvin Benson Cycle (CBC), **(C)** major carbohydrate (CHO) metabolism, **(D)** thioredoxin, **(E)** redox homeostasis, **(F)** photorespiration (PR), **(G)** nitrogen and sulfur metabolism (N/S) and **(H)** protein metabolism. The protein metabolism group is further divided in to three subgroups including synthesis, modification and degradation (from up to down). Data are the means of three to six biological replicates. The proteins high-lighted with pink background are validated Trx targets, while those with yellow background have been proposed to be redox-regulated proteins according to previous studies (Lindahl and Kieselbach, 2009). red = increased oxidation, blue = decreased oxidation. Raw data and statistics, see Supplemental Table S1.

While the oxidation states of these photosynthetic proteins stayed low for up to 2 h in the light, they surprisingly showed increasing re-oxidation after 6 h of illumination (Fig. 2, A, B and C). This indicates that after several hours of light exposure also oxidative processes come into play, leading to inactivation of a large set of photosynthetic proteins. Interestingly, CP12-1 and CP12-2, involved in regulatory-complex formation with phosphoribulokinase (PRK) and glyceraldehyde-3-phosphate dehydrogenase (GAPDH) in the CBC, showed specifically strong decreases in their oxidation levels after 10 min light, while there was no substantial re-oxidation at later time points, indicating both CP12 proteins to be less sensitive to light-dependent oxidation processes in the chloroplast.

To directly verify our redox proteomics results by an independent method, we selected three target proteins, namely plastidial fructose 1,6-bisphosphatase (CFBP), glyceraldehyde-3-phosphate dehydrogenase B (GAPB) and PRK, to be analyzed by protein electrophoretic mobility shift assay to assess light-dependent kinetics of their redox states using the same plant material taken for redox proteomics. The reduced thiols of proteins were alkylated using NEM, and the oxidized thiols of proteins were released by treating with DTT. The released thiols were further labeled with methoxypolyethlene glycol maleimide, which resulted in an increase of protein mass of the oxidized form, so that it became distinguishable from the reduced form during gel-electrophoresis. After immunoblotting, the intensity of the oxidized form was divided by the intensity sum of oxidized and reduced forms to yield the oxidation percentage of the respective protein. The gel blots in Supplemental Figure S2 show that all three proteins were fully oxidized in the dark (EN) yielding oxidation percentages of 100 % as shown in Figure 3. Compared to EN (dark), the oxidation percentages of CFBP and GAPB dropped down in the first 10 min to around 30% and continued to decrease within the next 2 h, while they showed subsequent re-oxidation after 6 h of light exposure (Supplemental Figure S2 and Fig. 3). The oxidation percentage of the PRK protein decreased very strongly already after 10 min of illumination down to levels that were hardly detectable, showing that this protein was very efficiently reduced by the dark-to-light transition (Supplemental Figure S2 and Fig. 3). These results obtained by gel-shift assays as an independent method are in confirmation with the redox proteomics data, with the light-dependent changes of the redox states of CFBP, GAPB and PRK being highly corresponding between the two different methods (compare Fig. 3 and Fig. 2B). Overall, these results validate our redox-proteomics data and the reliability of our experiment by an independent method, indicating that the redox-proteomics method we are using here is appropriate to determine light-dependent dynamics in protein redox states.

**Figure 3.**
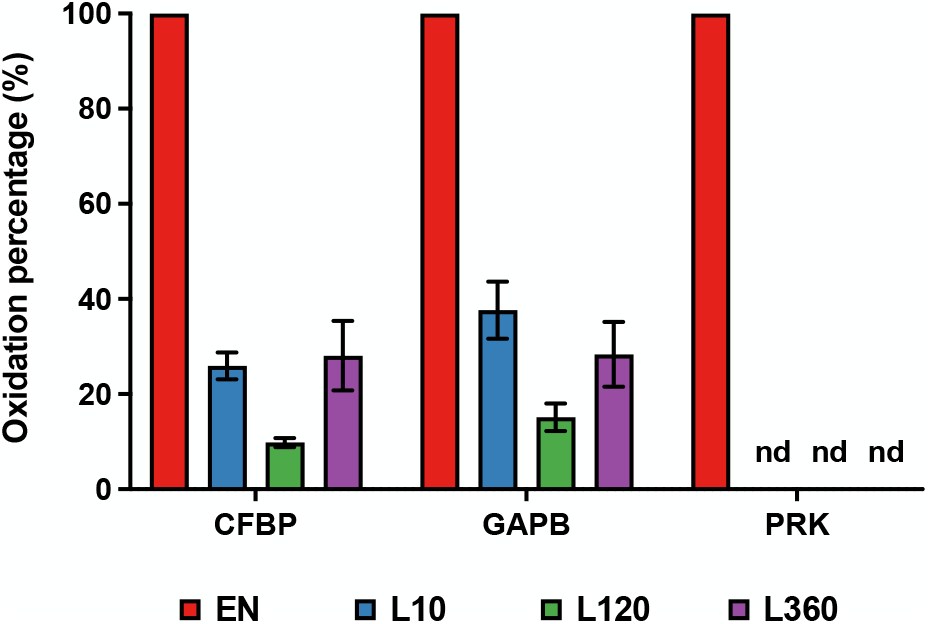
Validation of the redox proteomics results by analyzing oxidation percentages of CBC enzymes using protein electrophoretic mobility shift assays as an independent method. The protein oxidation percentages of chloroplastic fructose 1,6-bisphosphatase (CFBP), glyceraldehyde-3-phosphate dehydrogenase B (GAPB) and phosphoribulokinase (PRK) were analyzed via protein electrophoretic mobility shift assay as an independent method. For three independent biological replicates, the reduced thiols of proteins were alkylated using NEM, and the oxidized thiols of proteins were released by treating with DTT. The released thiols were further labeled with methoxypolyethlene glycol maleimide, which resulted in an increase of protein mass of the oxidized form that became distinguishable from the reduced form during gel electrophoresis. The oxidation percentages of CFBP, GAPB and PRK were calculated from the scanned blots shown in Supplemental Figure S2 by dividing the intensities of the bands reflecting the oxidized form by the sum of the intensities of the bands reflecting oxidized plus reduced forms of the respective protein. Results are means +/- SE (*n*=3). The symbol “nd” indicates “not detectable”.

### Light leads to a more complex pattern of redox changes in proteins involved in redox regulation, (photo)respiration, and protein metabolism

While protein redox changes are most likely linked to cellular Trx systems (Baumann and Juttner, 2002; Meyer et al., 2005; Geigenberger et al., 2017; Kang et al., 2019), light-dependent redox dynamics of Trx proteins were also analyzed (Fig. 2, D and E). Chloroplasts contain two different Trx systems, the Fdx/Trx system which is directly linked to light and proposed to activate photosynthetic target enzymes, and the NTRC system which is linked to NADPH and proposed as the major system to donate electrons to 2-Cys-Prx to convert H_2_O_2_ to H_2_O (Zaffagnini et al., 2019). Fdx-dependent chloroplast Trxs such as Trxs *f*1, *f*2, *m*1, *m*2, *m*4 and *x* showed a rapid decrease in their oxidation states within the first 10 min, followed by a re-increase after 120 and 360 min of light exposure (Fig. 2D), showing similar light-induced reduction-oxidation dynamics as their photosynthetic targets (Fig. 2B). This suggests *f*-, *m*- and *x*-types of Trxs to be subject to two opposing redox processes, leading to their reduction and re-oxidation in the light. While reduction is most-likely mediated by photo-reduced Fdx via FTR, acting within seconds to minutes of light exposure (Wang et al., 2014; Yoshida et al. 2022), the nature of the re-oxidation processes in the light is more obscure, but may be connected to 2-Cys peroxiredoxins (2-Cys Prxs; Lampl et al., 2022).

In comparison to this, both NADPH-dependent NTRC and Fdx-dependent Trx *y*2 showed an opposing pattern, being oxidized in the light, compared to the dark (Fig. 2D). NTRC (Kirchsteiger et al., 2009; Cejudo et al., 2012; Pérez-Ruiz et al., 2017) and Trx *y*2 (Collin et al., 2004; Shin et al., 2020) have been proposed in previous studies to act as efficient electron donors to 2-Cys Prx and Prx Q, respectively, to decompose H_2_O_2_ to H_2_O. Interaction of Trx *y*2 with 2-Cys Prx has also been demonstrated in a further study (Jurado-Flores et al., 2020), suggesting Trx *y*2 to be involved in re-oxidation processes related to the 2-Cys-Prx system. Interestingly, chloroplast 2-Cys Prxs and Prx-IIE showed similar light-dependent increases in their oxidation states, compared to NTRC and Trx *y*2 (Fig. 2, D and E). In addition to this, chloroplast superoxide dismutase 2 (CSD2) involved in antioxidative function showed an even stronger rise in its oxidation state upon illumination (Fig. 2E). The light-induced increase in oxidation of this set of proteins is most likely due to increased peroxide and ROS production during active photosynthetic processes in the chloroplast. This contrasts with other chloroplast antioxidative proteins, such as FSD1, MDAR6, GSH1 and APX4, and NAD(P)-dependent MDH, showing decreases in their oxidation states upon light exposure, similar to photosynthetic enzymes (Fig. 2E). This confirms previous studies, showing light-dependent reductive activation of NADP-MDH involved in the export of reducing equivalents from the chloroplast (Scheibe, 1991).

Thioredoxins are also residing outside the chloroplast, where they are reduced by NADP-dependent NTRA and NTRB (Reichheld et al., 2007; Bashandy et al., 2009; Cha et al., 2014). Our data show that cytosolic (Trxs *h*5 and *h*3) and mitochondrial (Trx *o*1) Trxs were oxidized in response to light, as well as NTRA (Fig. 2D). However, oxidation of the latter occurred only on a short-term basis (10 min light). Interestingly, with the exception of monodehydroascorbate reductase (MDAR1) and GDP-mannose 3,5-epimerase (GME), extra-plastidial glutathione peroxidases and reductases (GPX2, GPX6, GPX8 and ATGR1), superoxide dismutase (CSD1) and Prx-IIF were oxidized in the light (Fig. 2E), showing a similar pattern as extra-plastidial Trxs (Fig. 2D). Increased light-dependent oxidation of extra-plastidial Trxs and related peroxidases is probably due to increased ROS and H_2_O_2_ production during photosynthesis.

We also identified light-dependent changes in oxidation states in several proteins involved in photorespiration. Two peroxisomal enzymes, glycerate dehydrogenase (HPR) and glycolate oxidase (GLO1), were markedly reduced, while two mitochondrial glycine dehydrogenase proteins (GDH1 and GDH3) were oxidized during the day (Fig. 2F), indicating illumination might trigger differential redox regulation for photorespiratory processes in various subcellular compartments. When plants experienced a dark-to-light transition, many proteins involved in amino acid metabolism were also subject to redox changes (Fig. 2G). Notably, three plastidial enzymes showed strong redox changes during the day. The aspartate aminotransferase (ASP5) and the Fdx-dependent glutamate synthase 1 (GLU1) underwent strong reduction upon illumination, while the NADH-dependent glutamate synthase 1 (GLT1) showed an opposite redox pattern. In addition to this, we identified a proposed redox-regulated enzyme, cysteine synthase (OASB; Lindahl and Kieselbach, 2009), showing a strong reduction pattern during the day (Fig. 2G). It is worth noting that three sulfur-assimilation-related enzymes, ATP sulfurylase (ATPS1, ATPS2 and ATPS3), were significantly oxidized during the day (Fig. 2G), suggesting that also the sulfur metabolism is redox-regulated in response to light.

Interestingly, targets involved in plastidial protein metabolism also displayed clear redox changes during dark-to-light transition (Fig. 2H). These proteins were generally reduced in response to light, except a 50S ribosomal protein (RPL13) and two peptide methionine sulfoxide reductases (MSRB2 and MSR4). Notably, three of our identified targets including the chloroplastic elongation factor (CPEFG), MSR4 and the protease (CLPP5), were proposed to be redox-regulated targets in previous studies (Fig. 2H; Lindahl and Kieselbach, 2009). This may indicate that light is modulating plastidial protein homeostasis by redox regulation of a large set of target proteins in chloroplasts, confirming previous results showing that global translation is subject to redox-regulation in Arabidopsis (Moore et al., 2016) and yeast (Topf et al., 2018).

### Global changes of the redox proteome across mutants deficient in *m*-type Trxs, *f*-type Trxs or NTRC under various light conditions

Plastidial Trx proteins are crucial for light-dependent post-translational redox-regulation of photosynthetic metabolism (Collin et al., 2003; Geigenberger et al., 2017; Kang et al., 2019) and subject to reduction and oxidation in response to light-dependent processes (see above). To obtain more insights into the role of the plastidial thiol-redox system to regulate the cellular thiol-redox proteome, Arabidopsis mutants lacking parts of the Fdx-Trx or NADPH-NTRC systems were analyzed. We selected mutants lacking *f*-type (*trxf1f2*) and *m*-type Trxs (*trxm1m2*) as major and important parts of the chloroplast Fdx-Trx system, and the *ntrc* mutant lacking NADPH-dependent NTRC in the plastid. All three T-DNA insertion mutants have been extensively characterized in previous studies (Serrato et al., 2004; Naranjo et al., 2016a; Thormählen et al., 2017). For confirmation, expression levels of the respective genes were evaluated in these mutants, using real-time qPCR (Supplemental Fig. S3). As expected, Trx *f*1 and *f*2 signals were undetectable in the *trxf1f2* double mutant, and no Ntr*C* signal was detected in the *ntrc* mutant, indicating both lines to be null mutants. In the *trxm1m2* double mutant, no Trx *m*1 signal was detected, while the expression of Trx *m*2 was decreased down to approx. 60% of wild-type levels. Plants were grown in two different light conditions: In medium light (ML) as in the experiments described above, and in fluctuation light (FL) consisting of rapidly alternating high light (HL) and low light (LL) phases of 1 and 5 min, respectively. To waive potential diurnal effects and focus on investigating the roles of Trxs and NTRC in maintaining protein redox states, whole rosette leaves were harvested at six hours into the photoperiod for redox proteomics analyses.

Through the biotin-switch approach, we successfully identified 2220 proteins with redox-active Cys residues. After removing low-abundance targets (detected in less than 3 biological replicates) and subsequent statistical analyses (ANOVA with Dunnett’s test), 772 proteins were selected for the following analyses, which showed significant changes (P<0.05) in their oxidation levels when compared to the wild-type (Supplemental Table S4). Since the redox proteomics method used in this study does not provide absolute oxidation/reduction ratios of proteins (see above) we calculated the protein oxidation states of the mutants relative to the wild type. Data from a recent proteomics study indicate that this approach is not subject to substantial errors with respect to changes in protein abundance. Indeed, in constant ML or FL conditions, there were only minor changes in the overall abundance of proteins in trx*f*1, *trxm1m2* and *ntrc* mutants, relative to wild type (Dziubek et al., 2023). A comparison of our present data set (Supplemental Table S4) with those of Dziubek et al., (2023) pinpointed only 49 of the proteins that are relevant to the subsequent analyses and shown in the following data displays to reveal significant changes in protein abundance in the mutants relative to wild type (Supplemental Table S5). More crucially, as revealed in Supplemental Table S5, these proteins showed only very minor changes in their quantified levels (less than 3%) when mutants were compared to the wild type, indicating that changes in protein expression levels can be neglected as possible errors in our study.

The identified targets were grouped according to their subcellular localization and biological functions (Fig. 4, A and B). Most targets were localized in plastid (37% of total), cytosol (25% of total) and mitochondria (9% of total), while the rest (29% of total) distributes to other subcellular compartments (Figure 4A). As shown in Figure 4B, a large set of targets was associated to photosynthesis (13% of total), cellular respiration (5% of total), metabolic pathways (16% of total), redox homeostasis (5% of total) and RNA/protein processes (26% of total), while those remaining (36% of total) were allocated to the group of other cellular processes and unknown functions (Fig. 4B).

**Figure 4.**
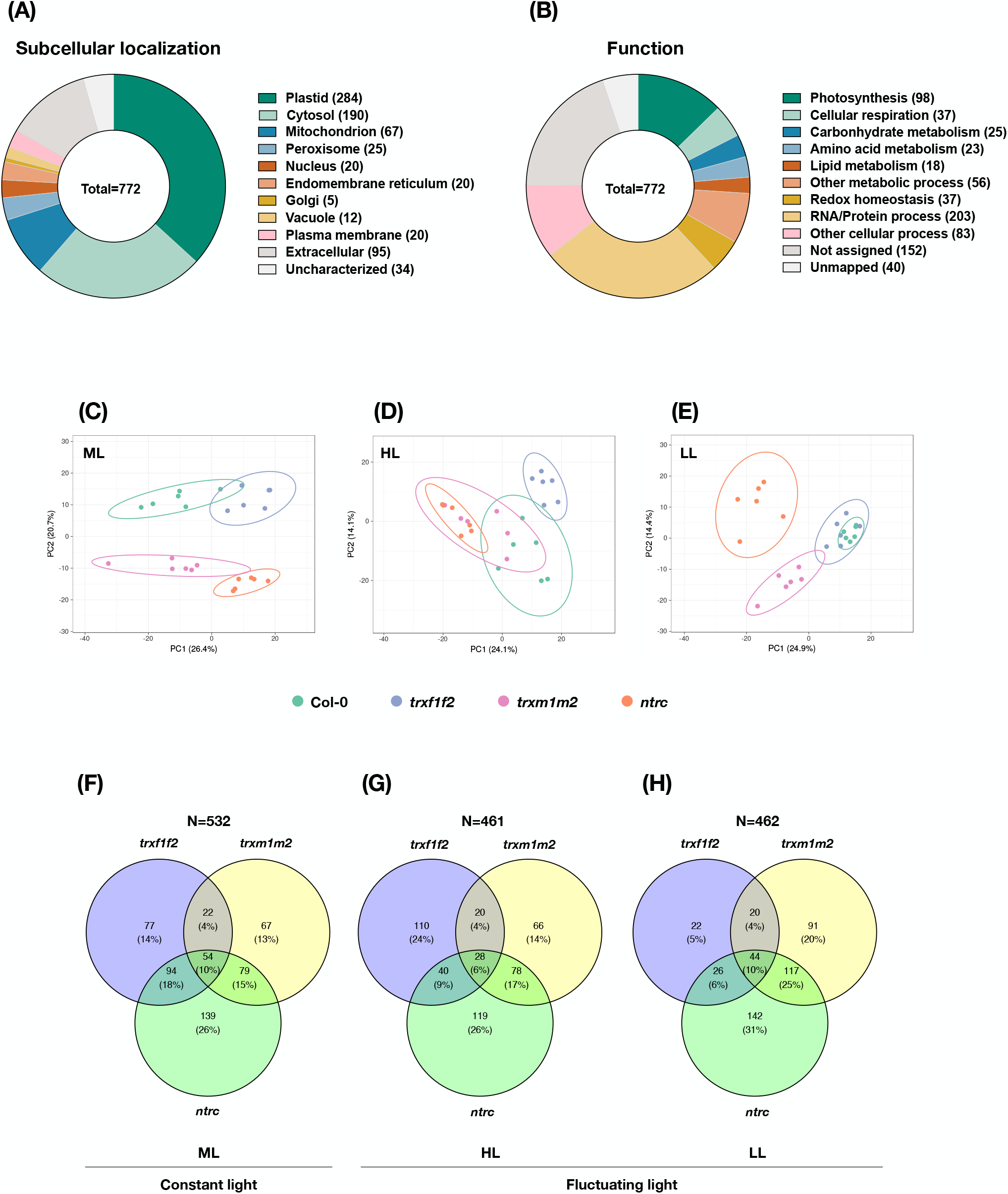
Dynamics in the landscape of protein oxidation changes across mutants lacking different parts of NTRC/Trx systems in constant and fluctuating light. Redox proteomics analyses of Arabidopsis *trxf1f2*, *trxm1m2* or *ntrc* mutants relative to the wild type (Col-0). Plants were grown in the same conditions as in Figures 1-3 (ML = medium light) or in fluctuating light (FL) with rapidly alternating high light (HL, 1 min) and low light (LL, 5 min) phases. Samples were taken 360 min into the photoperiod. **(A)** The subcellular localization of proteins showing significantly different oxidation states in the mutants relative to the wild type. **(B)** The functional categories of proteins showing significantly different oxidation states in the mutants relative to the wild type. **(C)** - **(E)** The principal component analysis displaying the distinctions of protein oxidation states between wild type (Col-0), *trxf1f2*, *trxm1m2* and *ntrc* mutants in ML or in HL or LL phases of FL. **(F)** - **(H)** Venn diagram highlighting the distribution of proteins subject to marked changes in their oxidation states in the respective mutants, compared to the wild type, during constant ML, as well as during HL or LL phases of FL. Data are based on three to six biological replicates.

We further performed PCA to visualize the effects of various light conditions on protein oxidation levels in different genotypes. In ML, the four genotypes formed four different clusters. The cluster of the *trxf1f2* double mutant was close to the wild type (Col-0) with a slight overlapping, while the clusters of *trxm1m2* and *ntrc* mutant lines stayed close to each other but were clearly deviated from the clusters of Col-0 and *trxf1f2* (Fig. 4C). When looking at the HL phases of FL, the clusters of *trxf1f2* and *ntrc* clearly deviated from the wild-type cluster, while the cluster of *trxm1m2* was largely overlapping with the clusters of Col-0 and *ntrc* (Fig. 4D). In the LL phases of FL, the *trxf1f2* double mutant clustered together with the wild type, while both *trxm1m2* and *ntrc* clustered differently and clearly deviated from the wild type (Fig. 4E). Taken together, these data indicate that *f*-type Trxs, *m*-type Trxs and NTRC have differential effects on the redoxome in ML, while in FL, *f*-type Trxs have a more specific effect on the redoxome in the HL phases, but *m*-type Trxs and NTRC in the LL phases.

Furthermore, we used Venn diagrams to visualize specific mutant effects on protein redox states in different light conditions. In ML, the *ntrc* mutants harbored 26% of identified proteins showing marked changes in their oxidation levels with respect to the wild type, suggesting a major role of NTRC in regulating global protein redox states, while the *trxf1f2* and *trxm1m2* double mutants comprised only 14% and 13% of identified proteins, respectively (Fig. 4F). Comparable amounts of targets resided in the overlapping region between *trxf1f2* and *ntrc* (18% of total) as well as *trxm1m2* and *ntrc* mutant lines (15% of total), indicating NTRC to influence the redox states of both *f* and *m*-type Trx target proteins, most likely in an indirect manner (Pérez-Ruiz et al., 2017). Nevertheless, only 4% of identified proteins were located in the overlapping region between *trxf1f2* and *trxm1m2* mutants (Fig. 4F), indicating the target specificities of *f* and *m*-type Trxs being rather distinct. Interestingly, there were 10% of identified proteins showing differential changes in their oxidation states with respect to the wild type in either of *trxf1f2*, *trxm1m2* or *ntrc* mutant lines. In FL, the *ntrc* mutant comprised the largest number of identified proteins showing significant changes in their oxidation levels relative to the wild type, in both HL (26% of total; Fig. 4G) and LL phases (31% of total; Fig. 4H). Interestingly, deficiencies of *f*-type Trxs led to larger changes in protein redox states (24% of total) in the HL phases (Fig. 4G) than in the LL phases (5% of total; Fig. 4H). This contrasts with deficiencies in *m*-type Trxs, which led to larger effects in the LL phases (20% of total) than in the HL phases (14% of total) of FL (Fig. 4H). The number of identified proteins residing in the overlapping regions between *ntrc* and *trxm1m2* were higher than in those between *ntrc* and *trxf1f2*, specifically in the LL phases (Fig. 4, G and H). This indicates a differential impact of *f*- and *m*-type Trxs on their target proteins in the HL and LL phases of FL, reflecting the different influence of NTRC on the redox states of these target proteins in the different light phases, which is mainly due to indirect effects (Fig. 4, G and H). Similar to ML conditions (Fig. 4F), in both HL (Fig. 4G) and LL phases of FL (Fig. 4H) only 4% of the identified proteins were located in the overlapping region between *trxf1f2* and *trxm1m2* mutants, indicating very distinct target specificities of *f-* and *m*-type Trxs independent of the light conditions.

### Deficiencies in *f*-type Trxs, *m*-type Trxs or NTRC differentially affect the redox proteome of photosynthesis and carbohydrate metabolism dependent on the light conditions

To further understand the contribution of plastidial thioredoxins (pTrxs) on the global redox proteome, we calculated the fold changes of protein oxidation levels in the mutant lines with respect to the wild type and categorized the targets showing significant difference (ANOVA with Dunnett’s test) into more detailed functional groups (Supplemental Table S4). In ML, deficiencies in *f-*, *m*-type Trxs or NTRC similarly led to a diverse pattern in the oxidation levels of proteins of the different photosystems (Fig. 5, A and B). When looking at the detailed changes of respective proteins, many PSI proteins became more oxidized, while several oxygen evolving enhancer proteins and PSII proteins were getting reduced in the *trxf1f2* mutants, compared to the wild type (Fig. 5, A and B). In comparison to the wild type, deficiency of NTRC led to a diverse redox pattern on PSI proteins, while the changes of PSII proteins in the *ntrc* mutant were similar to those in the *trxf1f2* mutant. In the *trxm1m2* double mutant, a large set of PSII proteins became more oxidized compared to the wild type, while the redox changes of PSI proteins were more diverse (Fig. 5, A and B). It is worth to note that deficiencies of *f-*, *m*-type Trxs or NTRC led to a mild increase in oxidation pattern in most electron carriers and ATP synthases except the cytochrome b6f proteins (PETA and PETC; Fig. 5C).

**Figure 5.**
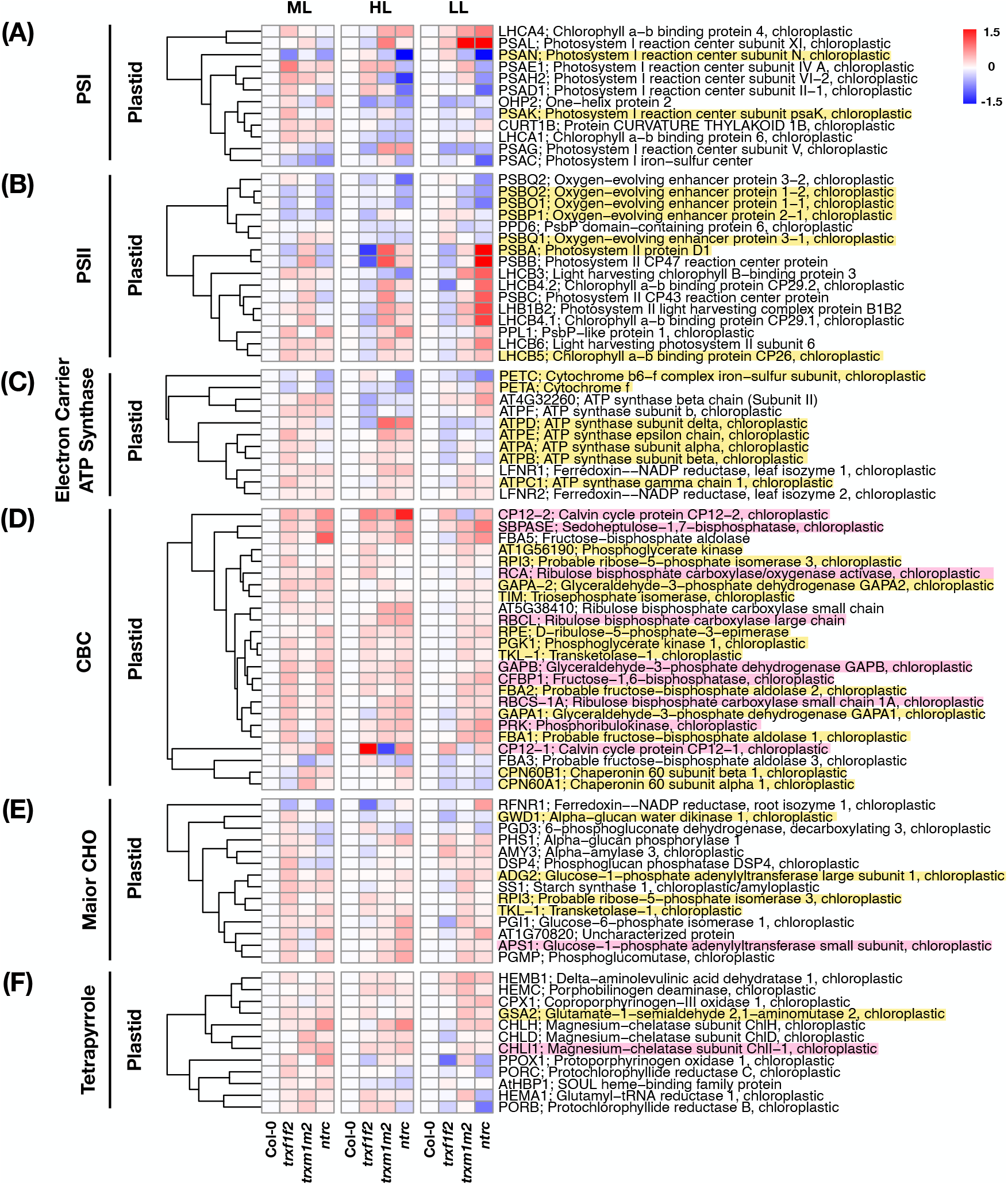
Protein oxidation changes in photosynthetic processes and carbohydrate metabolism across *trxf1f2*, *trxm1m2* and *ntrc* mutants in constant and fluctuating light. Heatmaps summarize the log_2_ fold changes of protein oxidation levels in Arabidopsis *trxf1f2*, *trxm1m2* or *ntrc* mutants relative to the wild type (Col-0). The proteins high-lighted with pink background are validated Trx targets, while those with yellow background have been proposed to be redox-regulated proteins according to previous studies (Lindahl and Kieselbach, 2009). Proteins were allocated to different groups of biological functions including **(A)** photosystem I (PSI), **(B)** photosystem II (PSII), **(C)** electron carrier and ATP synthase, **(D)** Calvin-Benson Cycle (CBC), **(E)** major carbohydrate (CHO) metabolism and **(F)** Tetrapyrrole. Plants were grown in the same conditions as indicated in the legend of Figure 4. Data are the means of three to six biological replicates. ML, medium light; FL, fluctuating light; HL, high-light phase of FL; LL, low-light phase of FL; red = increased oxidation, blue = decreased oxidation. Raw data and statistics, see Supplemental Table S4.

In contrast to this, during FL, the *trx* and *ntrc* mutant lines displayed distinct redox patterns in photosystem proteins. In the *trxf1f2* double mutant, the PSI proteins underwent diverse and minor redox changes (Fig. 5A), while the PSII proteins showed a general reduction pattern during the HL phases, compared to the wild type. Such reductive pattern was mitigated during the LL phases (Fig. 5B). Deficiency of *ntrc* led to marked reduction in most PSI proteins during either HL or LL phases. The redox patterns of PSI proteins in *trxm1m2* mutants were similar to those in *ntrc* mutants during the HL phases, while such reduction pattern was not maintained during the LL phases (Fig. 5A). When looking at the PSII group, the *ntrc* mutants showed a reduction pattern in oxygen-evolving proteins, but an oxidation pattern in other PSII reaction center proteins and chlorophyll-binding proteins, compared to the wild type. Such redox changes became more significant in the *ntrc* mutants during the LL phases (Fig. 5B). The redox patterns of PSII proteins in *trxm1m2* were similar to those in *ntrc* mutants. It is worth noting that the *trxf1f2* mutants exhibited a consistent reduction pattern on D1, CP47 and CP43 proteins (PSBA, PSBB and PSBC), while the *trxm1m2* mutants displayed opposite redox patterns on these three targets. This suggests *f* and *m*-type Trxs to be involved in an opposing manner in the assembly and repair of PSII. Indeed, it has been proposed that the *m*-type Trxs play a role in the biogenesis of PSII (Wang et al., 2013). Compromising *f*-type Trxs merely led to reduction of several ATP synthase proteins compared to the wild type (Fig. 5C). Nevertheless, deficiencies in *m*-type Trxs or NTRC similarly resulted in oxidation of most ATP synthase proteins during the HL phases, while such redox changes became less significant during the LL phases (Fig. 5C). Interestingly, in the *ntrc* mutants, the redox patterns of PETA and two ATP synthases (AT4G32260 and ATPF) appeared to be opposite between HL and LL phases (Fig. 5C). Taken together, the diverse redox changes of photosystem proteins in the different mutants indicate that different types of Trxs differentially affect the redox state of proteins of the photosynthetic light reactions.

Looking at enzymes of the CBC, deficiencies in Trxs *f*1/*f*2, Trxs *m*1/*m*2 or NTRC led to increased oxidation states of the respective proteins (Fig. 5D). This is in line with previous studies showing CBC enzymes to represent clear and confirmed targets of *f*- and *m*-type Trxs with different affinities, (Lindahl and Kieselbach, 2009; Yoshida et al., 2015; McFarlane et al., 2019; Yu et al., 2020), while the effect of NTRC on the redox state of these targets was shown to be indirect (Ojeda et al., 2017; Pérez-Ruiz et al., 2017). Interestingly, there were differences in the impact of these thiol-redox regulators on the redox-state of CBC targets depending on the light conditions. In ML, deficiencies of *f*-type Trxs or NTRC resulted in marked oxidation of almost all CBC proteins, while compromising *m*-type Trxs hardly affected the oxidation states of these targets (Fig. 5D). In FL, a different situation emerged. In the HL phases of FL, all three mutant lines displayed largely similar oxidation levels of CBC proteins, except a reduction in CP12-1 in the *trxm1m2* double mutant. When shifted to the LL phases, the redox states of most CBC enzymes remained oxidized in the *trxm1m2* and *ntrc* mutant lines, while those of the *trxf1f2* mutants showed a re-reduction to wild-type levels (Fig. 5D). While these results are in line with the generally accepted roles of Fdx-Trxs and NADPH-NTRC to modulate the reduction and hence the activation state of CBC enzymes (Michelet et al., 2013) they surprisingly show their different impacts depending on the light conditions. Specifically, our results indicate different impacts of *f*-type and *m*-type Trxs in reducing CBC enzymes in ML and FL conditions, with *m*-type Trxs playing a more important role in FL than in ML, and *f*-type Trxs being more important in ML and HL, rather than LL.

Next, we looked at the group of major CHO metabolism in the plastid. In ML, deficiencies of *f*-type Trxs led to a general oxidation of most CHO metabolism enzymes, while compromising *m*-type Trxs had only very minor effects on the redox states of these targets (Fig. 5E). Deficiency of NTRC resulted in diverse impacts on the redox states of CHO metabolism enzymes: A set of starch degradation enzymes was more reduced, while the CHO anabolic enzymes, including the well-known NTRC target, APS1, involved in starch synthesis, were more oxidized in the *ntrc* mutants compared to the wild type (Fig. 5E). In FL, no clear redox change of CHO metabolism enzymes was observed in the *trxf1f2* mutants. Nevertheless, during the LL phases, compromising *m*-type Trxs or NTRC led to a general oxidation in a large set of CHO metabolism enzymes (Fig. 5E). With respect to the redox regulation of CHO enzymes, this indicates *f*-type Trxs to play a major role in ML, while *m*- type Trxs appear to be more important in FL. Compromising NTRC led to increased oxidation of CHO enzymes in all light conditions, suggesting a more general role of NTRC in light regulation of carbohydrate metabolism. Notably, the RFNR1 was markedly reduced in the *trxf1f2* and *ntrc* mutant lines compared to the wild type in ML, and such reduction pattern was exacerbated only in the *trxf1f2* mutants during the HL phases of FL. Nevertheless, in the *ntrc* mutants, the redox state of RFNR1 was not altered during the HL phases but became more oxidized during the LL phases (Fig. 5E). Considering that the RFNR1 mediates the electron transfer between oxidative pentose phosphate pathway (OPPP) and downstream enzymes (Hanke et al., 2005), it is likely that the *f*-type Trxs and NTRC can regulate RFNR1 redox states to further modulate OPPP.

It has been reported that NTRC is involved in tetrapyrrole biosynthesis by regulating magnesium chelatase (Richter et al., 2013). Indeed, the target proteins of tetrapyrrole metabolism were more oxidized in the *ntrc* mutants compared to the wild type in ML, confirming the positive role of NTRC in chlorophyll biosynthesis (Fig. 5F). In addition, several targets of tetrapyrrole metabolism appeared to be more oxidized in the *trxf1f2* and *trxm1m2* mutants compared to the wild type (Fig. 5F), indicating both *f* and *m*-type Trxs also to participate in chlorophyll metabolism, as has been reported previously (Da et al., 2017; Wittmann et al., 2023). Increased oxidation of these proteins was maintained in the HL phases of FL, with some exceptions showing rather reduced status compared to the wild type (Figure 5F). Notably, the protoporphyrinogen oxidase (PPOX1) was dramatically reduced in the *trxf1f2* and *ntrc* mutant lines during the LL phases. Furthermore, the protochlorophylide reductases (PORB and PORC) and glutamyl-tRNA reductase (HEMA1) underwent reduction exclusively in the *ntrc* mutants during the LL phases, compared to WT (Fig. 5F).

### Deficiencies in *f*-type Trxs, *m*-type Trxs or NTRC affect the oxidation states of proteins involved in redox homeostasis, photorespiration, nitrogen and sulfur metabolism

We also evaluated the role of Trxs to catalyze redox changes in proteins involved in redox homeostasis. In ML, deficiencies of *f*-type Trxs or NTRC led to diverse redox changes in plastidial targets, while lack of *m*-type Trxs hardly changed the redox states of most targets (Fig. 6A) with the exception of superoxide dismutase [Fe] 1 (FSD1), which underwent marked oxidation in the *trxm1m2* mutant, but significant reduction in the *ntrc* mutant. Another superoxide dismutase [Cu-Zn] 2 (CSD2) and glutathione peroxidase (GPX1) were greatly oxidized in the *ntrc* mutant (Fig. 6A). Moreover, compromising the *m*/*f*-type Trxs or NTRC led to general reduction patterns in extra-plastidial targets including peroxiredoxins and enzymes of ascorbate-glutathione (AsA-GSH) cycle (Fig. 6A). Interestingly, there were changes in the redox pattern in the different phases of FL. In the HL phases, the different mutants led to similar changes as in ML, with the exception that in the *ntrc* mutant, CSD2 and GPX1 were less oxidized, while FSD1, Fdx-Trx reductase (FTRC), peroxiredoxins, and GSH S-transferase (DHA3) were more reduced, compared to the ML (Fig. 6A). This contrasts with the LL phases, where a set of targets including malate dehydrogenase (NADP- MDH), 2-Cys peroxiredoxin (BAS1), BAS1-like protein (2-Cys Prx B) and certain enzymes of AsA-GSH cycle (APX4 and MDAR6) showed strongly increased oxidation states in the *trxm1m2* and *ntrc* and mild increases in *trxf1f2* mutants, compared to HL phases of FL or ML (Fig. 6A). These redox changes of NADP-MDH are in line with a previous study using Arabidopsis mutants documenting that *m*-type Trxs and NTRC are involved in the activation of plastidial NADP-MDH *in vivo* (Thormählen et al., 2017). In contrast to *m*-type Trxs, NTRC is acting via an indirect mechanism, since it did not lead to a reduction of NADP- MDH via direct interaction *in-vitro* (Delgado-Requerey et al., 2023). Interestingly, chloroplast NAD-MDH was slightly reduced, indicating that the oxidation states of NAD and NADPH dependent MDHs located in the chloroplast responded differently after transfer from ML to FL. Furthermore, FTRC protein involved in Trx reduction was strongly reduced in all three mutant lines in FL, but not ML. With respect to the redox changes in extra-plastidial targets involved in redox homeostasis in FL, compared to ML, there was a clear tendency to increased oxidation states in most of the proteins in all three mutants. This was especially marked in the LL phases of FL (Fig. 6A).

**Figure 6.**
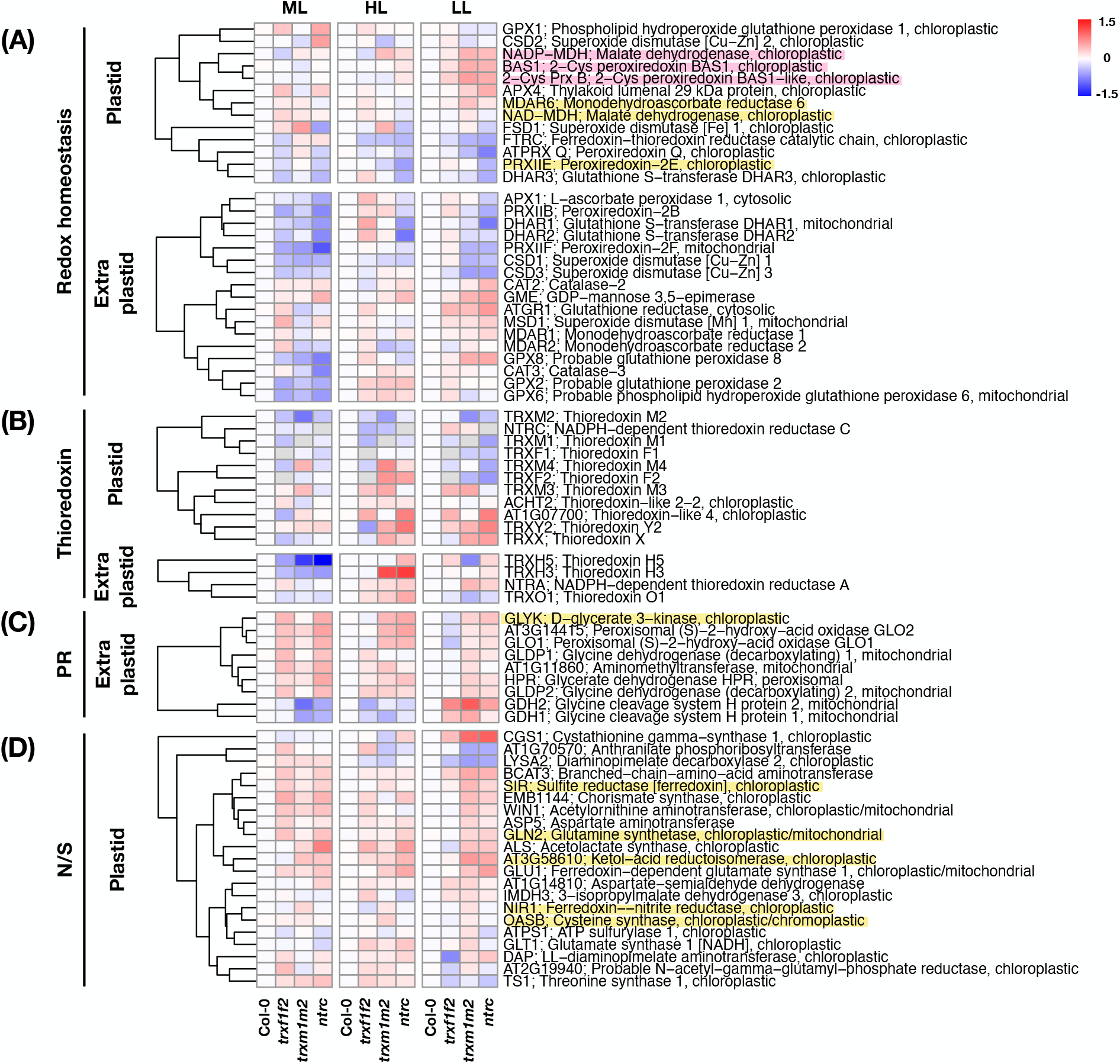
Protein oxidation changes in redox homeostasis, photorespiration and amino acid metabolism across *trxf1f2*, *trxm1m2* and *ntrc* mutants in constant and fluctuating light. Heatmaps summarize the log_2_ fold changes of protein oxidation levels in Arabidopsis *trxf1f2*, *trxm1m2* or *ntrc* mutants relative to the wild type (Col-0). The proteins high-lighted with pink background are validated Trx targets, while those with yellow background have been proposed to be redox-regulated proteins according to previous studies (Lindahl and Kieselbach, 2009). Proteins were allocated to different groups of biological functions including **(A)** redox homeostasis, **(B)** thioredoxin **(C)** photorespiration (PR) and **(D)** nitrogen and sulfur metabolism (N/S). Plants were grown in the same conditions as indicated in the legend to Figure 4. Data are the means of three to six biological replicates. ML, medium light; FL, fluctuating light; HL, high-light phase of FL; LL, low-light phase of FL; red = increased oxidation, blue = decreased oxidation. Raw data and statistics, see Supplemental Table S4.

There is evidence for cross talk between NTRC, 2-Cys Prx and Trxs *f* (Pérez-Ruiz et al., 2017) or Trxs *m* (Delgado-Requerey et al., 2023) in vivo. We thus investigated the redox changes of the other types of Trxs and Trx reductases in the selected mutant lines (Fig. 6B). In ML, deficiency of *f*-type Trxs elicited an increased reduction in Trx *m*2 and a reduction pattern in other Trxs, including plastidial Trxs *m*1, *m*4, *x* and NTRC as well as cytosolic Trxs *h*3 and *h*5, while the plastidial ACHT2 protein was more oxidized. Combined deficiencies of Trxs *m*1 and *m*2 instead led to increased oxidation of Trxs *m*3 and *m*4 in the plastid, while there was a decrease in the oxidation of extra-plastidial Trxs, specifically Trx *h*3 and *h*5 proteins in the cytosol. In response to NTRC deficiency, surprisingly, the oxidation of all other Trxs and Trx reductases was mitigated, except for cytosolic Trxs *h*3 and *h*5 showing strong decreases in their oxidation states (Fig. 6B). In FL, all three different mutants showed increased oxidation patterns in other Trxs and Trx reductases, when compared to ML (Fig. 6B). In the HL phases of FL, the rise in the oxidation states of those proteins was stronger than in LL phases, while *ntrc* and *trxm1m2* mutants showed more increased oxidation states than *trxf1f2* mutants, specifically with respect to the plastidial Trxs *f*2, *m*3, *m*4, y2, *x*, like-4, and ACHT2, and all extra-plastidial Trxs (Trxs *h*3, *h*5, *o*1 and NTRA). Taken together, deficiencies in *f*, *m*-type Trxs or NTRC led to differential effects in the redox states of other Trxs and Trx reductases dependent on the light conditions. While there were some mild decreases in oxidation in ML, oxidation states of most of those proteins were strongly increased in FL, specifically in the HL phases, with *ntrc* and *trxm1m2* having a much greater impact than *trxf1f2* mutants. This shows that in FL environments, NTRC and Trxs *m*1/*m*2 are specifically important to keep the other types of Trxs in a reduced state, also outside the plastid boundaries.

With respect to proteins of photorespiratory processes, all three mutants showed increased oxidation states, compared to wild type (Fig. 6C). In ML, *trxf1f2* and *ntrc* mutants showed the strongest effects on these photorespiratory targets, while in FL, specifically in the LL phases the impacts of *trxm1m2* and *ntrc* were more marked than those of *trxf1f2*. Interestingly, two glycine cleavage system proteins (GDH1 and GDH2) became less oxidized in *trxm1m2* and *ntrc* mutants in ML, while they showed a strong increase in oxidation in all three mutant lines in the LL phases of FL (Fig. 6C). Considering that most targets of photorespiration reside outside plastids, it is unlikely that the pTrxs or NTRC can directly modulate their redox states. Thus, the observed protein-redox changes are most likely the result of inter-organellar redox transfer by mechanisms such as the malate valve.

Following a similar pattern, almost all enzymes involved in plastid nitrogen and sulfur metabolism showed increased oxidation states in all mutants, compared to wild type (Fig. 6D). This involved several important enzymes such as sulfite reductase (SIR), glutamine synthetase (GLN2), Fdx-dependent glutamine synthase 1 (GLU1) and aspartate aminotransferase (ASP5), with SIR and GLN2 being suggested to be subject to redox-regulation in previous studies (Lindahl and Kieselbach, 2009). In ML, the impacts of *trxf1f2* and *ntrc* were stronger than those of *trxm1m2* mutants, while in the LL phases of FL, the impacts of *trxm1m2* became more dominant, while those of *trxf1f2* became diminished. This indicates *f*-, *m*-type Trxs and NTRC to be of general importance to keep the reduction states of enzymes of nitrogen and sulfur metabolism in a reduced state to optimize their activities in response to light. In this respect, the impacts of *f*-type and *m*-type Trxs were found to be different depending on the light conditions, being higher in ML and LL phases of FL, respectively.

### Deficiencies in *f*-type Trxs, *m*-type Trxs or NTRC affect the oxidation states of proteins involved in secondary metabolic pathways and protein homeostasis

We also looked at mutant effects on oxidation states of enzymes involved in other metabolic processes, such as lipid, nucleotide and secondary metabolism. Overall the three different mutants showed increased oxidation states of most of the proteins involved in these pathways, when compared to wild type (Supplemental Fig. S4). This included key enzymes of plastidial isoprenoid synthesis which have been suggested to be regulated by thiol/disulfide modulation in previous studies (Supplemental Fig. S4; Lindahl and Kieselbach, 2009) and shown to be involved in the methylerythritol 4-phosphate (MEP) pathway (i.e., 1- deoxy-D-xylulose 5-phosphate reductoisomerase, DXR; 4-hydroxy-3-methylbut-2-en-1-yl diphosphate synthase and reductase, ISPG and ISPH) and in the xanthophyll cycle (violaxanthin de-epoxidase, VDE1), where impacts of the different mutants on oxidation states were strongly dependent on light conditions. While in ML the oxidation states of these proteins were substantially increased, such oxidation patterns were mitigated in the HL phases, while there was a strong re-increase in the LL phases of FL (Supplemental Fig. S4). These dramatic changes in the oxidation pattern of these proteins between HL and LL phases of FL occurred within 1-4 min and were specifically marked in *trxm1m2* and *ntrc* mutants. This indicates that NTRC and Trxs *m*1/*m*2 are specifically important to balance the redox states of key enzymes of isoprenoid and zeaxanthin synthesis during rapidly altering HL and LL. The impacts of NTRC and Trxs *m*1/*m*2 were specifically strong with respect to VDE1, the key enzyme of zeaxanthin synthesis, which is crucial to decrease the photochemical efficiency of photosystem II by increasing heat dissipation via non-photochemical quenching (NPQ) to allow photoprotection during HL stress (Hieber et al., 2000). In confirmation to this, both, NTRC and Trxs *m*1/*m*2 were found to affect NPQ in the HL phases of FL (Thormählen et al., 2017).

Among the identified proteins, a surprisingly large set of targets involved RNA and protein processes in the plastid (Fig. 4B). We therefore had a closer look on genotypic changes in the oxidation states of these plastidial proteins (Supplemental Fig. S5). In ML, a large set of 30S and 50S ribosomal proteins and the elongation factors TUFA, emb2726 and CPEFG showed increases in their protein oxidation levels in all three mutant lines, compared to the wild type (Supplemental Fig. S5). This involved also proteins which have been suggested to be subject to thiol/disulfide regulation in previous studies (Supplemental Fig. S5; Lindahl and Kieselbach, 2009). Compared to ML, the oxidation levels of most of these proteins were found to be decreased in all three mutants in FL, specifically in the LL phases, where a set of 50S ribosomal proteins displayed a marked decrease in their oxidation states (Supplemental Fig. S5). Although deficiencies in *f*-, *m*-type Trxs or NTRC showed clear impacts on the oxidation levels of these ribosomal proteins, whether such redox changes affect protein metabolism requires further investigations.

## Discussion

Previous proteomics studies showed light variability with respect to diurnal changes (Uhrig et al., 2021) or fluctuating light (Niedermaier et al., 2020; Dziubek et al., 2023) to be associated with only minor changes in the overall abundance of proteins, pointing to the importance of thiol-disulfide modulation to regulate protein functions in response to light changes. However, in this context, our knowledge on the light-dependent dynamics of the plant redoxome is still scarce. In the present study, we performed biotin-switch based redox proteomics to systemically investigate the dynamics of light-dependent plant thiol-redox networks. By analyzing Arabidopsis plants at different time points into the photoperiod, we revealed illumination to lead to a marked increase in the reduction of a large set of proteins involved in photosynthetic processes during the first 10 min, followed by their partial re-oxidation after 2-6 h. Interestingly, *f*, *m* and *x*-type Trx proteins showed similar light-induced reduction-oxidation dynamics as their photosynthetic targets, while NTRC, 2-Cys-Prx and Trx *y*2 showed an opposing pattern, being more oxidized in the light, compared to the dark. This indicates the Trx/NTRC systems to be involved in, both, light-dependent reduction and re-oxidation dynamics. By analyzing Arabidopsis *trxf1f2*, *trxm1m2* and *ntrc* mutants, we found most protein targets to show increased oxidation states, compared to the wild type, suggesting their light-dependent decreases in oxidation states to be related to Trxs. Interestingly, *f*- and *m*-type Trxs were found to have different impacts on the thiol-redox proteome depending on the light conditions, with the impacts of Trxs *f*1/*f*2 to be higher in ML, while those of Trxs *m*1/*m*2 being increased in the LL-phases of FL. Compared to this, NTRC was found to have a strong impact in all light conditions. This indicates *f*-type Trxs, *m*-type Trxs and NTRC to be of general importance to keep the light-dependent thiol-redox proteome in a reduced state to optimize the functions of the constituent proteins, while they show different impacts depending on the light conditions.

### Light leads to reduction and re-oxidation dynamics of the plastid thiol-redox proteome

Our redox proteomics study shows that proteins revealing significant light- and Trx-dependent changes in their oxidation states were preferentially localized in the plastid (Fig. 1A and 4A) with photosynthesis being a major function (Fig. 1B and 4B). This is in agreement with a previous study, investigating the differential redox-modified proteins between regular growth light (GL) and FL in Arabidopsis, showing most proteins are localized in chloroplasts (Chen et al., 2022). Moreover, another redox proteomics study conducted in tobacco plants subjected to a very short-term dark-to-light transition also found that the identified proteins with redox-regulated and light-responsive properties are mainly localized in chloroplasts, even though the authors claimed their redox proteomics approach may not be suitable for proteins with multiple redox forms (Zimmer et al., 2021). Furthermore, a thioredoxome study in Chlamydomonas also found approx. 30% of 1188 Trx targets reside in chloroplasts (Pérez-Pérez et al., 2017). Taken together, light-dependent dynamics of the plant redox proteome are mainly localized to the chloroplast, where they are subject to the Trx systems linked to photo-reduced Fdx and NADPH-dependent NTRC.

When light-dependent changes in our redox proteomics data were analyzed in more detail, a large set of proteins involved in photosynthetic light reaction, CBC and carbohydrate metabolism showed marked decreases in their oxidation states within 10 min of illumination, compared to the dark (Fig. 2A). This observation is in line with the commonly accepted notion suggesting that illumination induces a rapid reductive signal pathway to activate chloroplast metabolism (Michelet et al., 2013). In confirmation to this, there was a rapid decrease in the oxidation states of chloroplast Trxs of *f*-type, *m*-type and *x*-type (Fig. 2D), which use electrons from photoreduced Fdx to reduce and activate plastidial targets. Interestingly, although such reduction patterns sustained within the next 2 hours of light exposure, oxidation levels increased again, returning almost to initial dark levels after six hours of illumination, indicating the occurrence of re-oxidation dynamics later in the photoperiod (Fig. 2, B and C; confirmed by an independent method in Fig. 3). This re-oxidation pattern was accompanied by increased oxidation states of *f*-, *m*- and *x*-type Trxs (Fig. 2D), which are probably attributable to an oxidation loop via 2-Cys-Prx (Zaffagnini et al. 2019). Interestingly, our present data show that 2-Cys-Prx, NTRC and Trx *y*2 are oxidized in the light, which supports this notion (Fig. 2, D and E).

Similar reduction and re-oxidation processes were previously documented when investigating the redox changes of NADP(H) couples during the dark-to-light transition, where the ratio of NADPH to the total pool of NADP+NADPH rapidly increased upon illumination, while recovering to the dark level after merely 3 min of light (Heber and Santarius, 1965; Dietz and Hell, 2015). Moreover, a recent study using biosensors to monitor cellular redox signals revealed that the Fdx-mediated reductive signals interact with the 2- Cys Prxs-mediated oxidative signals to fine-tune photosynthetic processes (Lampl et al., 2022). In fact, deficiencies of 2-Cys Prxs were reported to facilitate the reduction of certain CBC enzymes (Pérez-Ruiz et al., 2017). Moreover, co-incubating oxidized 2-Cys Prx with plastidial Trxs can effectively inactivate CFBP and MDH activities (Vaseghi et al., 2018). In this context, the light-dependent increase in H_2_O_2_ may account for the substantial oxidation of 2-Cys Prxs and its electron donor, NTRC, during the day (Fig. 2, D and E), as both components need to serve as electron sinks to maintain the oxidative signal transduction. Moreover, oxidation of NTRC in the light may also be due to the reoxidation pattern of the NADPH/NADP redox couple after 3 min of illumination (Heber and Santarius, 1965; Dietz and Hell, 2015). The finding that NTRC is oxidized in the light is in line with its proposed mechanism to act indirectly on plastidial targets via 2-Cys Prx (Pérez-Ruiz et al., 2017), rather than by their direct reduction (Ojeda et al., 2017). Interestingly, also Trx *y*2 has been proposed in previous studies to act as efficient electron donor to 2-Cys Prx (Collin et al., 2004; Jurado-Flores et al., 2020; Shin et al., 2020), while it was shown to be inefficient to directly reduce CBC proteins (Collin et al., 2004; Yoshida et al., 2015). The reason for this unexpected specificity of Trx *y*2 remains to be determined. In summary, our redox proteomics data reveal re-oxidation dynamics in the light, with the reduction and re-oxidation network being linked to the Fdx-FTR and NTRC-2-Cys-Prx systems, respectively, while plants use this redox network to fine-tune photosynthetic processes during the day. Interestingly, such 2-Cys Prxs-mediated protein oxidation is also characterized in human cell lines (Stöcker et al., 2018), indicating oxidative signal transduction to be ubiquitous among different organisms.

### NTRC as well as *f*- and *m*-type Trxs play major roles to reduce the proteins of plastid carbon metabolism with their individual impacts depending on the light conditions

Our redox proteomics data show that in constant ML, almost all CBC enzymes and proteins of related pathways displayed increased oxidation states in the *trxf1f2* and *ntrc* mutants relative to wild type (Fig. 5D), reinforcing the notion that, both, *f*-type Trxs and NTRC are involved in the redox-activation of CBC enzymes under normal light conditions *in vivo*, although the effect of the latter was found to be indirect (Yoshida et al., 2015; Yoshida and Hisabori, 2016; Guinea Diaz et al., 2020; Thormählen et al. 2015). In contrast to this, the *trxm1m2* mutant showed only a minor impact on CBC protein oxidation states in ML, but a stronger impact in FL, indicating *m*-type Trxs to be more important to regulate the redox states of CBC enzymes in FL than in the ML environments (Fig. 5D). Indeed, several studies proposed a specific importance of *m*-type Trxs, but not *f*-type Trxs, in photosynthetic acclimation to FL and LL environments (Thormählen et al., 2017; Da et al., 2018; Okegawa and Motohashi, 2020). In contrast to this, deficiency of NTRC strongly increased the oxidation levels of CBC proteins and proteins of related pathways, both, in ML and FL (Fig. 5D). The wide oxidation effect of NTRC deficiency on plastidial targets is most likely attributable to decreased provision of electrons to 2-Cys Prx leading to increased oxidation of plastidial Trxs (Vaseghi et al., 2018; Cejudo et al., 2019). To sum up, these results indicate plants to flexibly adopt different Trx systems to optimize redox regulation of plastidial targets in acclimation to different light conditions.

### NTRC controls the oxidation levels of proteins of photosynthetic light reactions

Our data show that deficiency in NTRC also affects the oxidation levels of proteins of the photosynthetic machinery, which is more evident in FL than in ML, where PSI proteins became more strongly reduced, while PSII proteins showed markedly increased oxidation levels (Fig. 5, A and B). It has been proposed that deficiency of NTRC restricts the electron donation to PSI (Naranjo et al., 2016b), but this seems to have minor effects on the oxidation levels of PSI proteins in ML (Fig. 5A). In contrast, NTRC deficiency led to a marked increase in reduction in PSI proteins in FL (Fig. 5A), which is most likely attributable to decreased electron transfer from PSI to CBC due to an inhibition of the later. Indeed, previous studies show that in FL, *ntrc* mutants display higher acceptor side limitation of PSI (Y(NA)) than the wild type (Nikkanen et al., 2018). Subsequently, this over-reduction of PSI will promote ROS generation, which may ultimately lead to increased oxidation of ROS-sensitive PSII proteins (Fig. 5B). Collectively, the results suggest that NTRC is an important hub in controlling the redox homeostasis of light reactions, especially in FL. This is in line with *ntrc* mutants showing a decreased photosynthetic performance, specifically in FL (Thormählen et al., 2017). In contrast, *trxf1f2* and *trxm1m2* mutants exhibited only mild and inconsistent oxidation changes of photosystem proteins, compared to wild type (Fig. 5, A and B). The relevance of the redox changes of photosystem proteins to determine the photosynthetic performance of these mutants requires further investigations.

### NTRC/Trx systems regulate rapid changes in the oxidation state of proteins involved in isoprenoid synthesis during alternating HL and LL phases of FL

Our results show that the oxidation states of key enzymes of plastidial isoprenoid synthesis, which have been suggested to be regulated by thiol/disulfide modulation in previous studies, were strongly affected in *trxf1f2*, *trxm1m2* or *ntrc* mutants, depending on the light conditions (Supplemental Fig. S3). There were specifically strong effects on VDE1. In ML, the oxidation states of VDE1 in all three mutant lines were only slightly altered, compared to wild type. This observation is in line with a previous study, showing that deficiency of NTRC does not significantly affect VDE redox state in regular light conditions (Naranjo et al., 2016b). This differs in plants growing in FL environments. Here VDE1 was subject to a dramatic decrease in oxidation level in *trxm1m2* and *ntrc* mutants, specifically during the HL phases of FL (1 min), while there was a marked increase in its oxidation status during the subsequent LL phases (5 min; Supplemental Fig. S3). VDE proteins are the key enzymes of the xanthophyll cycle, which is responsible for dissipating excess light energy (Havaux et al., 2007; Fernández-Marín et al., 2021). An *in vitro* assay suggested that VDE remains active only when it is completely oxidized (Yamamoto and Kamite, 1972; Simionato et al., 2015). The de-epoxidation of violaxanthin is usually more active during HL as plants need to dissipate excess light energy. In this context, the decreased VDE1 oxidation in *trxf1f2, trxm1m2* and *ntrc* mutants suggests the interference of violaxanthin de-epoxidation in the HL phases of FL. Collectively, the results indicate that the NTRC-Trx system operates to regulate VDE1 redox states when plant experience FL.

### The influence of NTRC or Trxs *m*1/*m*2 on the redox states of other chloroplast Trxs is specifically strong during rapid fluctuations in light intensities

Our redox proteomics data showed that 6-h into the photoperiod in ML, Trxs *f*1/*f*2, Trxs *m*1/*m*2 and NTRC deficiencies had only minor effects on the oxidation levels of other pTrxs (Fig. 6B). Compared to this, the dysregulation of the redox balance in other pTrxs became clearer in FL, specifically in *ntrc* und *trxm1m2* mutants, showing an increased oxidation pattern in most of these proteins in the HL periods (Fig. 6B). This is in line with previous studies showing increased oxidation states of Trxs *f*1 and *f*2 in the *ntrc* mutant, compared to wild-type, as a short-term response during dark-light transitions (Pérez-Ruiz et al., 2017). Accordingly, we hypothesize a role of NTRC and Trxs *m*1/*m*2 to balance the redox state of other types of Trxs to cope with short-term light fluctuations. This is in line with the hypothesis suggesting NTRC to indirectly modulate the redox states of other pTrxs, especially the *f*-type Trxs, via its role in balancing the 2-Cys Prx redox state (Cejudo et al., 2019). Mechanistically speaking, the operation of NTRC substantially provides reducing equivalents to 2-Cys Prxs and therefore minimizes the drainage of reducing equivalents from the pools of other types of Trxs to 2-Cys Prxs (Vaseghi et al., 2018; Cejudo et al., 2019). Our data indicate that in this context, the impact of NTRC is stronger during short-term fluctuations in light intensity than in long-term constant light.

### Proteins of nitrogen and sulfur metabolism are linked to light-responsive redox regulation via NTRC-Trx systems

The marked changes in the oxidation states of GLU1 and GLT1 upon illumination led us to assume that nitrogen metabolism is under the thiol-redox control in response to light (Fig. 2G). It has been proposed in previous studies that the glutamine synthetase (GS)/glutamate synthase (GOGAT) plays a central role in leaf nitrogen assimilation. GLU1 as the major GOGAT enzyme is predominately expressed in leaf tissues, and it accounts for 95% of the GOGAT activity (Somerville and Ogren, 1980; Coschigano et al., 1998; Coruzzi, 2003). GLU1 mainly uses the electrons from Fdx to catalyze glutamate synthesis (Suzuki and Knaff, 2005). Its activity and protein accumulation are enhanced during the day (Coschigano et al., 1998; Schjoerring et al., 2006). In fact, we detected less oxidized form of GLU1 protein upon illumination (Fig. 2G). In the context, the accumulating GLU1 proteins is subject to a strong reduction during the day. This is in line with previous studies, showing DTT treatment or addition of pTrxs to activate Fdx-dependent GOGAT isolated from spinach chloroplasts (Lichter and Haberlein, 1998). The reduction of GLU1 may therefore lead to increased activity of the GS/GOGAT cycle during the day. Interestingly, the NADH-dependent GOGAT, GLT1, showed a marked oxidation pattern upon illumination. Such oxidation may explain why GLT1 maintains in a very low level of leaf GOGAT activity (Somerville and Ogren, 1980; Coschigano et al., 1998; Coruzzi, 2003). Our data also indicate redox-control of aspartate synthesis. The oxidized level of ASP5 markedly decreased during the day (Fig. 2G), indicating its strong reduction. However, this probably does not change aspartate synthesis since previous studies analyzing a missense mutant of ASP5 shows that interfering ASP5 activity does not change the levels of aspartate and asparagine (Miesak and Coruzzi, 2002). Nevertheless, our results suggest ASP5 harbors redox-active Cys residues, and its redox state is regulated by light. Furthermore, the oxidation levels of GLU1 and ASP5 moderately increased in mutants deficient in *f*-, *m*-type Trxs or NTRC, confirming their regulation by the Trx/NTRC systems (Fig. 6D), which may explain changes in amino acid accumulation (Thormählen et al., 2015).

Our data also show interesting changes in the oxidation states of proteins involved in sulfur metabolism. Upon illumination, the oxidation levels of three ATP sulfurylase (ATPS) proteins were significantly increased (Fig. 2G), which is in line with previous study showing ATPS proteins to be targets of Trxs (Marchand et al., 2006). Fdx-dependent sulfite reductase (SIR1), a further enzyme of S-assimilation which has been confirmed as Trx target in previous studies (Lindahl and Kieselbach, 2009), showed increased oxidation levels in t*rxf1f2*, *trxm1m2* and *ntrc* mutants, compared to wild type (Fig. 6D). Taken together, our redox proteomics data provide additional evidence that nitrogen and sulfur metabolism are associated with NTRC/Trx-mediated redox regulation *in vivo*.

### Protein metabolism is subject to redox regulation via NTRC/Trx systems

Our redox proteomics data show that a large set of proteins involved in different processes of protein metabolism were subject to changes in their oxidation levels in response to illumination (Fig. 1B and 4B), indicating redox regulation to be also operational in protein homeostasis. This is in line with two recent articles suggesting thiol-based redox switches to be operational at each step of translation and to play a major role in controlling protein homeostasis (Moore et al., 2016; Topf et al., 2018). In the redoxome conducted in yeast, the authors proposed that ribosomal proteins can serve as sensors to monitor cellular redox states to modulate translation processes in response to environmental changes (Topf et al., 2018). This suggests that the decrease in oxidation levels of plastidial ribosomal proteins during dark-to-light transition may act as an additional “kick-off” signal to activate photosynthetic processes. There is also evidence that the redox states of these plastidial ribosomal proteins are under the control of the NTRC/Trx systems, since their oxidation levels were markedly changed in mutants deficient in *f*-, *m*-type Trx or NTRC in ML (Supplemental Fig. S5).

The underlying mechanisms by which redox modifications of ribosomal proteins modulate translational processes are still unclear. Interestingly, in our redox-proteomics study we identified a well-known chloroplastic elongation factor (CPEFG), which was characterized by markedly decreased oxidation levels upon illumination, while its oxidation levels increased in *trxf1f2*, *trxm1m2* and *ntrc* mutants relative to wild type in either ML or FL (Fig. 2H and Supplemental Fig. S5). This indicates the redox state of CPEFG to be controlled by NTRC/Trx systems in response to light. In confirmation to this, the homologue of CPEFG in the cyanobacterium *Synechocystis* was found to be subject to redox regulation by the NTR-Trx system in previous studies. The reduced form of CPEFG can actively facilitate translation processes (Kojima et al., 2009). Furthermore, deficiency of CPEFG in Arabidopsis was found to strongly delay the accumulation of plastid proteins (LHCP, D1 and CP22) including Rubisco subunits, resulting in an *albino* phenotype at early developmental stages (Albrecht et al., 2006). Taken together, we propose that Trx/NTRC-dependent reduction of CPEFG upon illumination will facilitate the translation of plastid-encoded transcripts to optimize protein homeostasis inside the chloroplast.

### Conclusion remark

In this study we used redox proteomics to systematically investigate the dynamics of the thiol-redox network in plants in response to temporal changes in light availability and across genotypes lacking different parts of NTRC/Trx systems. We found light to lead to reduction and re-oxidation dynamics of photosynthetic proteins linked to the Fdx/Trx (*f*, *m* and *x*-type Trxs) and the NTRC/2-Cys-Prx systems (including Trx *y*2), respectively, which showed opposite changes in their light-responsive redox patterns. While deficiencies in *f*-type Trxs, *m*-type Trxs or NTRC were mainly associated with increased oxidation states of photosynthetic proteins, their impacts differed in different light environments, with NTRC and Trxs *f*1/*f*2 being important to keep proteins in a reduced state in constant light, while NTRC and Trxs *m*1/*m*2 being indispensable to balance oxidation/reduction-dynamics of proteins during rapid alterations in light intensity in FL environments.

## Materials and Methods

### Plant material and growth conditions

The wild-type Arabidopsis plant, Columbia-0 (Col-0), and the well-characterized T-DNA insertion mutant lines *trxf1f2* (SALK_128365/GK-020E05-013161; Naranjo et al., 2016a), *trxm1m2* double mutants (SALK_123570/SALK_130686; Thormählen et al., 2017) and *ntrc* single mutant (SALK_012208, Serrato et al., 2004), were used for the following analyses. All plants were grown in a growth chamber equipped with LED light. The light intensity was set as 150 μmol photons m^-2^ s^-1^ with a 12-h-dark/12-h-light regime and the temperature was set as 22°C. After three weeks, half of the plants were shifted to fluctuating light (FL) where they were repeatedly exposed to 1 min high light (HL; 550 μmol photons m^-2^ s^-1^) and 5 min low light (LL; 50 μmol photons m^-2^ s^-1^) in the same dark-to-light regime, while the other half remained in the initial constant medium light (ML) conditions. The plants were then grown for another week in the respective light conditions, before whole rosette leaves were sampled six hours into the photoperiod by shock freezing in liquid nitrogen. In FL, leaf samples were taken in HL and LL periods separately.

#### RNA extraction, reverse transcription and real-time quantitative PCR

The leaf material was frozen in liquid nitrogen and ground to fine powder. Approximate 50 mg of leaf powder was used for RNA extraction by implementing RNAzol reagent (Sigma-Aldrich). The RNA concentration was determined using NanoDrop spectrophotometer (ND- 2000, ThermoFisher Scientific), and 500 ng of total RNA was used to synthesize cDNA using iScript reverse transcription kit (Biorad). The cDNA sample was diluted 20 times with nuclease-free water, and 5 μL of diluted cDNA was implemented for real-time quantitative PCR using the SYBG reagent (Biorad). The PCR reaction was performed in the thermocycler (C1000 Touch^TM^ Thermal Cycler; Biorad). The detailed protocol can be found in the previous publication (Hou et al., 2021), and the primer pairs used in the real-time quantitative PCR were listed below: TRXf1_qFW (5’-cgatgatctggttgcagcg-3’), TRXf1_qRv (5’- ctggttcatccggaagcag-3’), TRXf2_qFW (5’-tgtaaccaagacaacaagcca-3’), TRXf2_qRv (5’- cggtcacttcctttactacct-3’), TRXm1_qFW (5’-taacactgatgagtctcctgcaa-3’), TRXm1_qRv (5’- gatgctggttgctaaagtgtctt-3’), TRXm2_qFW (5’-tgaagctcaggaaactactaccgat-3’), TRXm2_qRV (5’-cagtgtaatgctgtgctagatcg-3’), NtrC_qFw (5’-tgaagatgaagaaagagtaccgag-3’), NtrC_qRv (5’- ggtgtcctcatttattggcct-3’).

#### Protein extraction and biotin-switch labeling method

The biotin-switch assays were performed according to published methods with several modifications (Jaffrey and Snyder, 2001; Liu et al., 2014). Briefly, 50 mg of finely pulverized leaf sample was re-suspended in 350 μL of extraction buffer (20 mM Tris-HCl, pH 7.0, 5 mM EDTA, 100 mM NaCl, 6M Urea) containing 50 mM NEM, and the whole extract was incubated for 30 min at 25°C with mild shaking in the dark to alkylate free thiols. The leaf debris were removed by centrifugation at 20,000 x *g* for 10 min at 4°C, and the supernatant was mixed with 4 volumes of absolute acetone to precipitate protein. The protein pellet was recovered, cleaned up and dehydrated as mentioned above. The protein pellet was further re-suspended in 200 μL of extraction buffer containing 100 mM dithiothreitol (DTT) followed by incubation for 30 min at 37°C with mild shaking in the dark to release oxidized thiols. Protein extract was subject to acetone precipitation, cleaned up and dehydrated as above. The protein pellet was re-suspended in 200 μL of extraction buffer, and the protein concentration was determined using 660 nm protein reagent (Pierce). Approximate 80 μg of total protein was used for biotin labeling in the presence of 0.4 mM thiol-reactive N-[6-(Biotinamido) hexyl]-3’-(2’-pyridyldithio) propionamide (biotin-HPDP; Cayman Chemical). The sample was then incubated for 60 min at 25°C with mild shaking in the dark followed by acetone precipitation and cleanup. The protein pellet was first re-suspended in 50 μL of extraction buffer and further diluted by adding 450 μL of binding buffer (20 mM Tris-HCl, pH 7.0; 5 mM EDTA; 100 mM NaCl). The biotinylated protein extract was incubated with 100 μL Streptavidin resin (Invitrogen) for 60 min at 25°C with mild shaking in the dark. The resin was recovered by centrifugation at 2000 xg for 1 min at 4°C. The protein-bound resin was washed three times with 500 μL of binding buffer and twice with 500 μL of 20 mM ammonium bicarbonate solution. The resin was incubated in 200 μL of binding buffer containing 100 mM DTT for 30 min at 37°C with mild shaking in the dark to elute bound proteins. The eluted proteins were subject to acetone precipitation, cleaned up and dehydrated.

#### Mass spectrometry

In Solution Tryptic Digest and LC-MS/MS Analysis was basically performed as described by (Hammel et al., 2018). Proteins were resuspended in 8M urea / ammonia bicarbonate buffer, digested using LysC and trypsin and desalted using home-made C18-STAGE tips. Finally, the peptides were resuspended in a solution of 2% acetonitrile, 2% formic acid. The LC-MS/MS system (Eksigent nanoLC 425 coupled to a TripleTOF 6600, ABSciex) was operated in μ-flow mode using a 25 μ-emitter needle in the ESI source. Peptides were separated by reversed phase (Triart C18, 5 μm particles, 0.5 mm × 5 mm as trapping column and Triart C18, 3 μm particles, 300 μm × 150 mm as analytical column, YMC) using a flow rate of 4 μl/min and a gradient ramping from 1 to 5% HPLC buffer B (buffer A 2% acetonitrile, 0.1% formic acid; buffer B 90% acetonitrile, 0.1% formic acid) within 5 min, to 35% buffer B in 73 min and to 50% buffer B in 2 min, followed by wash and equilibration steps. The mass spectrometer was operated in data-dependent analysis with one MS1 spectrum (350 to 1,250 m/z, 250 ms) and 35 triggered MS/MS scans in high sensitivity mode (110 to 1,600 m/z, 50 ms) for >2 times charged ions, resulting in a total cycle time of 2,050 ms. Fragmented precursors were excluded for 10 s and precursors with a response below 150 cps were excluded completely from MS/MS analysis.

#### Mass spectrometry data analyses

Analysis of MS raw data was performed using MaxQuant version 1.6.0.1 using default settings with minor changes (Cox and Mann, 2008). Library generation for peptide spectrum matching was based on *Arabidopsis thaliana* (UniProt reference proteome UP0000065489) including chloroplast and mitochondrial proteins. Oxidation of methionine and acetylation of the N-termini were considered as peptide modifications. Maximal missed cleavages were set to 3 and peptide length to 6 amino acids, the maximal mass to 6,000 Da. Thresholds for peptide spectrum matching and protein identification were controlled for a false discovery rate (FDR) of 0.01. To account for the stochastic effect of data-dependent acquisition, match between runs (MBR) was used and to minimize the variation between different samples MaxQuant label-free quantification (LFQ) intensity values were reported. Raw data were deposited at PRIDE proteome exchange with identifier PXD043914. For further statistical analysis the identified proteins been detected in less than three biological replicates were considered as low-abundance targets, and omitted from the following data processing. The majority proteins IDs were used for the following annotation. The proteins IDs were converted to AGI locus codes using the online mapping tool provided by UniProt (https://www.uniprot.org/id-mapping/). The AGI locus codes were used to obtain gene names using the online tool, EnsemblPlants. The subcellular localizations of proteins were yielded from the SUBA4 database (Hooper et al., 2017). The biological functions of proteins were grouped using MapMan according to Araport and TAIR databases. The unsupervised cluster analyses were conducted using MapMan as well. The PCA was performed using the online tool, ClustVis (Metsalu and Vilo, 2015). The heatmaps, Venn diagrams and statistical analyses were performed using the R program. The other plots were graphed using the Graphpad Prism version 9.

### Validation of the results by an independent method

Redox proteomics results were validated by electrophoretic mobility shift assays as an independent method. The protein redox states of CFBPase, GAPDH and PRK were analyzed via protein electrophoretic mobility shift assay as described previously with several modifications (Muthuramalingam et al., 2010; Nikkanen et al., 2016). In brief, 20 mg of ground leaf powder was re-suspended in 200 μL of 10% (w/v) Tricholoroacetic acid solution followed by incubation on ice for 20 min. The supernatant was removed by centrifugation at 20,000 xg for 10 min at 4°C, and the protein pellet was washed twice with 1 mL of 80% (v/v) acetone prepared in 50 mM Tris-HCl (pH7.0). The protein pellet was dehydrated and then re-suspended in 200 μL of extraction buffer (100 mM Tris-HCl, pH7.5, 1 mM EDTA, 2% [w/v] SDS, 6 M urea, proteinase inhibitor cocktail) containing 50 mM N-ethylmaleimide (NEM) followed by incubation for 30 min at 25°C with mild shaking in the dark to alkylate free thiols. The supernatant was recovered via centrifugation at 20,000 xg for 5 min at 4°C. Equal amount of 20% (w/v) Trichloroacetic acid solution was applied into the protein extract, and the whole mixture was incubated on ice for 20 min. The protein pellet was recovered, cleaned up and dehydrated as mentioned above. The protein pellet was further re-suspended in 200 μL of extraction buffer containing 100 mM dithiothreitol (DTT) followed by incubation for 30 min at 37°C with mild shaking in the dark to release oxidized thiols. Equal amount of 20% (w/v) Tricholoroacetic acid solution was applied into the protein extract to stop the reaction, and the whole mixture was incubated on ice for 20 min. The protein pellet was recovered, cleaned up and dehydrated as mentioned above. The protein pellet was next re-suspended in 100 μL of extraction buffer containing 10 mM of methoxypolyethlene glycol maleimide (Mal-PEG, 5 kDa, Sigma-Aldrich_63187) followed by incubation for 60 min at 27°C with mild shaking in the dark to label the reduced thiols. Two microliter of 1 M DTT was applied to stop the reaction, and 500 μL of absolute acetone was applied into the protein extract to precipitate protein and get rid of excess Mal-PEG. The protein pellet was recovered, cleaned up and dehydrated as mentioned above. The protein pellet was solubilized in 2-time-strength Laemmli buffer (Laemmli, 1970). The protein samples were analyzed by SDS-PAGE and Western blot. CFBP was detected using a commercial FBPase1 antibody (Agrisera; AS194319), while GAPB and PRK were detected using antibodies as described previously (Teh et al., 2023). Theoretically, the mass shift of the protein, in which a single reduced disulphide was labeled with Mal-PEG, is 10 kDa. Nevertheless, due to the hydration of PEG, the actual mass shift could become larger, up to 22 kDa (Makmura et al., 2001; Peled-Zehavi et al., 2010).

## Supplemental Data

**Supplemental Figure S1. The procedures of samples preparation and redox proteomics.** (A) Arabidopsis plants were grown under light intensity of 150 μmol photons m^-2^ s^-1^ with a 12-h-dark/12-h-light regime for 4 weeks. To monitor a time-course in the light, whole rosette leaves were sampled by shock-freezing in liquid nitrogen at end of night (EN) and 10, 120 and 360 min into the photoperiod. (B) The leaf samples were used for protein extraction in the presence of NEM to block the free thiol residues followed by DTT treatment to reduce the oxidized thiols. The released thiols were labeled with redox-active biotin, and the protein extract was subject to affinity purification using a streptavidin column. The bound proteins were eluted via incubating with DTT.

**Supplemental Figure S2. Validation of the redox proteomics results by analyzing oxidized and reduced forms of CBC enzymes using protein electrophoretic mobility shift assays as an independent method.** The protein oxidation percentages of chloroplastic fructose 1,6-bisphosphatase (CFBP), glyceraldehyde-3-phosphate dehydrogenase B (GAPB) and phosphoribulokinase (PRK) were analyzed via protein electrophoretic mobility shift assay as an independent method. For three independent biological replicates, the reduced thiols of proteins were alkylated using NEM, and the oxidized thiols of proteins were released by treating with DTT. The released thiols were further labeled with methoxypolyethlene glycol maleimide, which resulted in an increase of protein mass of the oxidized form that became distinguishable from the reduced form during gel electrophoresis. The oxidized and reduced forms of the proteins are indicated on the immunoblots. The immunosignals marked “ox” and “”re” represent the oxidized and reduced forms of proteins, respectively, while those marked with asterisk represent the signals of non-specific binding. The scanned blots were used to calculate oxidation percentages of CFBP, GAPB and PRK shown in Figure 3.

**Supplemental Figure S3. Molecular characterization of *trxf1f2*, *trxm1m2* and *ntrc* mutant lines.** The expression levels of Trx *f*1, *f*2, *m*1, *m*2 and Ntr *C* were detected using real-time quantitative PCR with gene-specific primers. The gene expression was quantified using 2^−ΔΔCt^ method. The symbol “nd” indicates “not detected”

**Supplemental Figure S4. Protein oxidation changes in lipid, nucleotide and secondary metabolism across *trxf1f2*, *trxm1m2* and *ntrc* mutants in constant and fluctuating light.** Heatmaps summarize the log_2_ fold changes of protein oxidation levels in Arabidopsis *trxf1f2*, *trxm1m2* or *ntrc* mutants relative to the wild type (Col-0). The targets with yellow background are proposed to be redox-regulated proteins according to previous studies (Lindahl and Kieselbach, 2009). Proteins were allocated to different groups of biological functions including lipid, nucleotide and secondary metabolism. Plants were grown in the same conditions as indicated in the legend to Figure 4. Data are the means of three to six biological replicates. ML, medium light; FL, fluctuating light; HL, high-light phase of FL; LL, low-light phase of FL; red = increased oxidation, blue = decreased oxidation. Raw data and statistics, see Supplemental Table S4.

**Supplemental Figure S5. Protein oxidation changes in protein metabolism across *trxf1f2*, *trxm1m2* and *ntrc* mutants in constant and fluctuating light.** Heatmaps summarize the log_2_ fold changes of protein oxidation levels in Arabidopsis *trxf1f2*, *trxm1m2* or *ntrc* mutants relative to the wild type (Col-0). The targets with yellow background are proposed to be redox-regulated proteins according to previous studies (Lindahl and Kieselbach, 2009). Proteins were allocated to different groups of biological functions including protein synthesis, modification and degradation. Plants were grown in the same conditions as indicated in the legend to Figure 4. Data are the means of three to six biological replicates. ML, medium light; FL, fluctuating light; HL, high-light phase of FL; LL, low-light phase of FL; red = increased oxidation, blue = decreased oxidation. Raw data and statistics, see Supplemental Table S4.

**Supplemental Table S1.** Proteins showing statistically significant changes in oxidation states in response to illumination compared to the dark.

**Supplemental Table S2.** Proteins with light-dependent changes in oxidation levels that have been reported to be also subject to diurnal changes with respect to protein abundance. By comparing the list of proteins identified to be subject to light-dependent changes in oxidation levels in Supplemental Table S1 with the published data set of Uhrig et al. (2021), we found that only 16 out of 319 proteins were also reported to be subject to diurnal changes in overall protein levels.

**Supplemental Table S3.** Proteins displaying significant redox changes in all three time points of the photoperiod compared to the dark.

**Supplemental Table S4.** Proteins showing significant redox changes in Arabidopsis *trxf1f2*, *trxm1m2* or *ntrc* mutants with respect to the wild type (Col-0) during ML and FL.

**Supplemental Table S5.** Comparison of our present data set (Supplemental Table S4) with those of Dziubek et al., (2023) in ML and FL pinpoints 49 of the proteins that are relevant to our analyses and shown in Figures 5, 6, S4 and S5 to reveal significant changes in protein abundance. These proteins showed only very minor changes in their quantified levels (less than 3%) when mutants were compared to the wild type (Col-0), indicating that changes in protein expression levels can be neglected as possible errors in our study.

## Funding

This work was supported by the Deutsche Forschungsgemeinschaft (TRR175).

## Acknowledgements

We are grateful to all green house staff at Faculty of Biology in Ludwig-Maximilians-University Munich for taking care of Arabidopsis plants.

## Literature cited

Albrecht V, Ingenfeld A, Apel K (2006) Characterization of the snowy cotyledon 1 mutant of Arabidopsis thaliana: The impact of chloroplast elongation factor G on chloroplast development and plant vitality. Plant Mol Biol 60: 507–518

Alkhalfioui F, Renard M, Montrichard F (2007) Unique properties of NADP- thioredoxin reductase C in legumes. J Exp Bot. doi: 10.1093/jxb/erl248

Arsova B, Hoja U, Wimmelbacher M, Greiner E, Üstün Ş, Melzer M, Petersen K, Lein W, Börnke F (2010) Plastidial thioredoxin z interacts with two fructokinase-like proteins in a thiol-dependent manner: Evidence for an essential role in chloroplast development in Arabidopsis and Nicotiana benthamiana. Plant Cell 22: 1498–1515

Balmer Y, Koller A, Del Val G, Manieri W, Schürmann P, Buchanan BB (2003) Proteomics gives insight into the regulatory function of chloroplast thioredoxins. Proc Natl Acad Sci U S A 100: 370–375

Balmer Y, Vensel WH, Hurkman WJ, Buchanan BB (2006) Thioredoxin Target Proteins in Chloroplast Thylakoid Membranes. Antioxid Redox Signal 8: 1829– 1834

Balmer Y, Vensel WH, Tanaka CK, Hurkman WJ, Gelhaye E, Rouhier N, Jacquot JP, Manieri W, Schürmann P, Droux M, et al (2004a) Thioredoxin links redox to the regulation of fundamental processes of plant mitochondria. Proc Natl Acad Sci U S A 101: 2642–2647

Balmer Y, Vensel WH, Tanaka CK, Hurkman WJ, Gelhaye E, Rouhier N, Jacquot J-P, Manieri W, Schurmann P, Droux M, et al (2004b) Thioredoxin links redox to the regulation of fundamental processes of plant mitochondria. Proceedings of the National Academy of Sciences 101: 2642–2647

Bartsch S, Monnet J, Selbach K, Quigley F, Gray J, Von Wettstein D, Reinbothe S, Reinbothe C (2008) Three thioredoxin targets in the inner envelope membrane of chloroplasts function in protein import and chlorophyll metabolism. Proc Natl Acad Sci U S A 105: 4933–4938

Bashandy T, Taconnat L, Renou JP, Meyer Y, Reichheld JP (2009) Accumulation of flavonoids in an ntra ntrb mutant leads to tolerance to UV-C. Mol Plant 2: 249–258

Bohrer AS, Massot V, Innocenti G, Reichheld JP, Issakidis-Bourguet E, Vanacker H (2012) New insights into the reduction systems of plastidial thioredoxins point out the unique properties of thioredoxin z from Arabidopsis. J Exp Bot 63: 6315–6323

Brandes HK, Larimer FW, Geck MK, Stringer CD, Schurmann P, Hartman FC (1993) Direct identification of the primary nucleophile of thioredoxin f. Journal of Biological Chemistry 268: 18411–18414

Carrillo LR, Froehlich JE, Cruz JA, Savage LJ, Kramer DM (2016) Multi-level regulation of the chloroplast ATP synthase: The chloroplast NADPH thioredoxin reductase C (NTRC) is required for redox modulation specifically under low irradiance. The Plant Journal 87: 654–663

Cejudo FJ, Ferrández J, Cano B, Puerto-Galán L, Guinea M (2012) The function of the NADPH thioredoxin reductase C-2-Cys peroxiredoxin system in plastid redox regulation and signalling. FEBS Lett 586: 2974–2980

Cejudo FJ, González MC, Pérez-Ruiz JM (2021) Redox regulation of chloroplast metabolism. Plant Physiol 186: 9–21

Cejudo FJ, Ojeda V, Delgado-Requerey V, González M, Pérez-Ruiz JM (2019) Chloroplast redox regulatory mechanisms in plant adaptation to light and darkness. Front Plant Sci 10: 1–11

Cha JY, Kim JY, Jung IJ, Kim MR, Melencion A, Alam SS, Yun DJ, Lee SY, Kim MG, Kim WY (2014) NADPH-dependent thioredoxin reductase A (NTRA) confers elevated tolerance to oxidative stress and drought. Plant Physiology and Biochemistry 80: 184–191

Chen Q, Xiao Y, Ming Y, Peng R, Hu J, Wang H, Jin H (2022) Quantitative proteomics reveals redox-based functional regulation of photosynthesis under fluctuating light in plants. J Integr Plant Biol 64: 2168–2186

Collin V, Issakidis-Bourguet E, Marchand C, Hirasawa M, Lancelin JM, Knaff DB, Miginiac-Maslow M (2003) The Arabidopsis plastidial thioredoxins. New functions and new insights into specificity. Journal of Biological Chemistry 278: 23747–23752

Collin V, Lamkemeyer P, Miginiac-Maslow M, Hirasawa M, Knaff DB, Dietz KJ, Issakidis-Bourguet E (2004) Characterization of plastidial thioredoxins from arabidopsis belonging to the new y-type. Plant Physiol 136: 4088–4095

Coruzzi GM (2003) Primary N-assimilation into Amino Acids in Arabidopsis. Arabidopsis Book 2: e0010

Coschigano KT, Melo-Oliveira R, Lim J, Coruzzi GM (1998) Arabidopsis gls mutants and distinct Fd-GOGAT genes: Implications for photorespiration and primary nitrogen assimilation. Plant Cell 10: 741–752

Cox J, Mann M (2008) MaxQuant enables high peptide identification rates, individualized p.p.b.-range mass accuracies and proteome-wide protein quantification. Nat Biotechnol 26: 1367–1372

Cremers CM, Jakob U (2013) Oxidant sensing by reversible disulfide bond formation. Journal of Biological Chemistry 288: 26489–26496

Da Q, Sun T, Wang M, Jin H, Li M, Feng D, Wang J, Wang H Bin, Liu B (2018) M-type thioredoxins are involved in the xanthophyll cycle and proton motive force to alter NPQ under low-light conditions in Arabidopsis. Plant Cell Rep 37: 279–291

Da Q, Wang P, Wang M, Sun T, Jin H, Liu B, Wang J, Grimm B, Wang H-B (2017) Thioredoxin and NADPH-dependent thioredoxin reductase C regulation of Tetrapyrrole Biosynthesis. Plant Physiol 175: 652–666

Delgado-Requerey V, Cejudo FJ, González M-C (2023) The Functional Relationship between NADPH Thioredoxin Reductase C, 2-Cys Peroxiredoxins, and m-Type Thioredoxins in the Regulation of Calvin–Benson Cycle and Malate-Valve Enzymes in Arabidopsis. Antioxidants 12: 1041

Dietz KJ, Hell R (2015) Thiol switches in redox regulation of chloroplasts: Balancing redox state, metabolism and oxidative stress. Biol Chem 396: 483–494

Dziubek D, Poeker L, Siemiątkowska B, Graf A, Marino G, Alseekh S, Arrivault S, Fernie AR, Armbruster U, Geigenberger P (2023) NTRC and thioredoxins *m* 1/ *m* 2 underpin the light acclimation of plants on proteome and metabolome levels. Plant Physiol. doi: 10.1093/plphys/kiad535

Fernández-Marín B, Roach T, Verhoeven A, García-Plazaola JI (2021) Shedding light on the dark side of xanthophyll cycles. New Phytologist 230: 1336–1344

Geigenberger P, Thormählen I, Daloso DM, Fernie AR (2017) The Unprecedented Versatility of the Plant Thioredoxin System. Trends Plant Sci 22: 249–262

Goyer A, Haslekås C, Miginiac-Maslow M, Klein U, Le Marechal P, Jacquot JP, Decottignies P (2002) Isolation and characterization of a thioredoxin-dependent peroxidase from Chlamydomonas reinhardtii. Eur J Biochem 269: 272–282

Guinea Diaz M, Nikkanen L, Himanen K, Toivola J, Rintamäki E (2020) Two chloroplast thioredoxin systems differentially modulate photosynthesis in Arabidopsis depending on light intensity and leaf age. Plant Journal 104: 718–734

Hall M, Mata-Cabana A, Åkerlund HE, Florencio FJ, Schröder WP, Lindahl M, Kieselbach T (2010) Thioredoxin targets of the plant chloroplast lumen and their implications for plastid function. Proteomics 10: 987–1001

Hammel A, Zimmer D, Sommer F, Mühlhaus T, Schroda M (2018) Absolute Quantification of Major Photosynthetic Protein Complexes in Chlamydomonas reinhardtii Using Quantification Concatamers (QconCATs). Front Plant Sci. doi: 10.3389/fpls.2018.01265

Hanke GT, Okutani S, Satomi Y, Takao T, Suzuki A, Hase T (2005) Multiple iso-proteins of FNR in Arabidopsis: Evidence for different contributions to chloroplast function and nitrogen assimilation. Plant Cell Environ 28: 1146–1157

Havaux M, Dall’Osto L, Bassi R (2007) Zeaxanthin Has Enhanced Antioxidant Capacity with Respect to All Other Xanthophylls in Arabidopsis Leaves and Functions Independent of Binding to PSII Antennae. Plant Physiol 145: 1506–1520

Heber UW, Santarius KA (1965) Compartmentation and reduction of pyridine nucleotides in relation to photosynthesis. Biochim Biophys Acta 109: 390–408

Hieber AD, Bugos RC, Yamamoto HY (2000) Plant lipocalins: violaxanthin de-epoxidase and zeaxanthin epoxidase. Biochimica et Biophysica Acta (BBA) - Protein Structure and Molecular Enzymology 1482: 84–91

Holmgren A (1995) Thioredoxin structure and mechanism: conformational changes on oxidation of the active-site sulfhydryls to a disulfide. Structure 3: 239–243

Hooper CM, Castleden IR, Tanz SK, Aryamanesh N, Millar AH (2017) SUBA4: the interactive data analysis centre for Arabidopsis subcellular protein locations. Nucleic Acids Res 45: D1064–D1074

Hou LY, Lehmann M, Geigenberger P (2021) Thioredoxin h2 and o1 show different subcellular localizations and redox-active functions, and are extrachloroplastic factors influencing photosynthetic performance in fluctuating light. Antioxidants. doi: 10.3390/antiox10050705

Jacquot JP, Eklund H, Rouhier N, Schürmann P (2009) Structural and evolutionary aspects of thioredoxin reductases in photosynthetic organisms. Trends Plant Sci 14: 336–343

Jaffrey SR, Snyder SH (2001) The Biotin Switch Method for the Detection of S - Nitrosylated Proteins. Science’s STKE. doi: 10.1126/stke.2001.86.pl1

Jurado-Flores A, Delgado-Requerey V, Gálvez-Ramírez A, Puerto-Galán L, Pérez-Ruiz JM, Cejudo FJ (2020) Exploring the functional relationship between ytype thioredoxins and 2-cys peroxiredoxins in arabidopsis chloroplasts. Antioxidants 9: 1–18

Kang Z, Qin T, Zhao Z (2019) Thioredoxins and thioredoxin reductase in chloroplasts: A review. Gene 706: 32–42

Kirchsteiger K, Pulido P, Gonzálezlez M, Cejudo FJ (2009) NADPH Thioredoxin reductase C controls the redox status of chloroplast 2-Cys peroxiredoxins in arabidopsis thaliana. Mol Plant 2: 298–307

Kojima K, Motohashi K, Morota T, Oshita M, Hisabori T, Hayashi H, Nishiyama Y (2009) Regulation of translation by the Redox State of Elongation factor G in the cyanobacterium Synechocystis sp. PCC 6803. Journal of Biological Chemistry 284: 18685–18691

Laemmli UK (1970) Cleavage of structural proteins during the assembly of the head of bacteriophage T4. Nature 227: 680–685

Lamkemeyer P, Laxa M, Collin V, Li W, Finkemeier I, Schöttler MA, Holtkamp V, Tognetti VB, Issakidis-Bourguet E, Kandlbinder A, et al (2006) Peroxiredoxin Q of Arabidopsis thaliana is attached to the thylakoids and functions in context of photosynthesis. Plant Journal 45: 968–981

Lampl N, Lev R, Nissan I, Gilad G, Hipsch M, Rosenwasser S (2022) Systematic monitoring of 2-Cys peroxiredoxin-derived redox signals unveiled its role in attenuating carbon assimilation rate. Proc Natl Acad Sci U S A 119: 1–10

Lemaire SD, Guillont B, Le Maréchal P, Keryer E, Miginiac-Maslow M, Decottignies P (2004) New thioredoxin targets in the unicellular photosynthetic eukaryote Chlamydomonas reinhardtii. Proc Natl Acad Sci U S A 101: 7475–7480

Lepistö A, Pakula E, Toivola J, Krieger-Liszkay A, Vignols F, Rintamäki E (2013) Deletion of chloroplast NADPH-dependent thioredoxin reductase results in inability to regulate starch synthesis and causes stunted growth under short-day photoperiods. J Exp Bot 64: 3843–3854

Lichter A, Haberlein I (1998) A light-dependent redox signal participates in the regulation of ammonia fixation in chloroplasts of higher plants - Ferredoxin: Glutamate synthase is a thioredoxin-dependent enzyme. J Plant Physiol 153: 83–90

Lindahl M, Florencio FJ (2003) Thioredoxin-linked processes in cyanobacteria are as numerous as in chloroplasts, but targets are different. Proc Natl Acad Sci U S A 100:16107–16112

Lindahl M, Kieselbach T (2009) Disulphide proteomes and interactions with thioredoxin on the track towards understanding redox regulation in chloroplasts and cyanobacteria. J Proteomics 72: 416–438

Liu P, Zhang H, Wang H, Xia Y (2014) Identification of redox-sensitive cysteines in the arabidopsis proteome using OxiTRAQ, a quantitative redox proteomics method. Proteomics 14: 750–762

Löwe O, Rezende F, Heidler J, Wittig I, Helfinger V, Brandes RP, Schröder K (2019) BIAM switch assay coupled to mass spectrometry identifies novel redox targets of NADPH oxidase 4. Redox Biol 21: 101125

Makmura L, Hamann M, Areopagita A, Furuta S, Muñoz A, Momand J (2001) Development of a Sensitive Assay to Detect Reversibly Oxidized Protein Cysteine Sulfhydryl Groups. Antioxid Redox Signal 3: 1105–1118

Marchand C, Le Maréchal P, Meyer Y, Decottignies P (2006) Comparative proteomic approaches for the isolation of proteins interacting with thioredoxin. Proteomics 6: 6528–6537

Marchand CH, Vanacker H, Collin V, Issakidis-Bourguet E, Le Maréchal P, Decottignies P (2010) Thioredoxin targets in Arabidopsis roots. Proteomics 10: 2418–2428

McFarlane CR, Shah NR, Kabasakal B V., Echeverria B, Cotton CAR, Bubeck D, Murray JW (2019) Structural basis of light-induced redox regulation in the Calvin– Benson cycle in cyanobacteria. Proceedings of the National Academy of Sciences 116: 20984–20990

Metsalu T, Vilo J (2015) ClustVis: a web tool for visualizing clustering of multivariate data using Principal Component Analysis and heatmap. Nucleic Acids Res 43: W566–W570

Meyer Y, Belin C, Delorme-Hinoux V, Reichheld J-P, Riondet C (2012) Thioredoxin and glutaredoxin systems in plants: Molecular mechanisms, crosstalks, and functional significance. Antioxid Redox Signal 17: 1124–1160

Michalska J, Zauber H, Buchanan BB, Cejudo FJ, Geigenberger P (2009) NTRC links built-in thioredoxin to light and sucrose in regulating starch synthesis in chloroplasts and amyloplasts. Proceedings of the National Academy of Sciences 106: 9908–9913

Michelet L, Lemaire SD, Marchand CH, Fermani S, Sparla F, Morisse S, Pérez-Pérez ME, Trost P, Zaffagnini M, Danon A, et al (2013) Redox regulation of the Calvin– Benson cycle: something old, something new. Front Plant Sci 4: 1–21

Miesak BH, Coruzzi GM (2002) Molecular and physiological analysis of Arabidopsis mutants defective in cytosolic or chloroplastic aspartate aminotransferase. Plant Physiol 129: 650–660

Montrichard F, Alkhalfioui F, Yano H, Vensel WH, Hurkman WJ, Buchanan BB (2009) Thioredoxin targets in plants: The first 30 years. J Proteomics 72: 452–474

Moore M, Gossmann N, Dietz KJ (2016) Redox Regulation of Cytosolic Translation in Plants. Trends Plant Sci 21: 388–397

Muthuramalingam M, Dietz K-J, Ströher E (2010) Thiol–Disulfide Redox Proteomics in Plant Research. NIH Public Access 639: 1–14

Naranjo B, Diaz-Espejo A, Lindahl M, Cejudo FJ (2016a) Type-f thioredoxins have a role in the short-term activation of carbon metabolism and their loss affects growth under short-day conditions in Arabidopsis thaliana. J Exp Bot 67: 1951–1964

Naranjo B, Mignée C, Krieger-Liszkay A, Hornero-Méndez D, Gallardo-Guerrero L, Cejudo FJ, Lindahl M (2016b) The chloroplast NADPH thioredoxin reductase C, NTRC, controls non-photochemical quenching of light energy and photosynthetic electron transport in Arabidopsis. Plant Cell Environ 39: 804–822

Navrot N, Collin V, Gualberto J, Gelhaye E, Hirasawa M, Rey P, Knaff DB, Issakidis E, Jacquot JP, Rouhier N (2006) Plant glutathione peroxidases are functional peroxiredoxins distributed in several subcellular compartments and regulated during biotic and abiotic stresses. Plant Physiol 142: 1364–1379

Niedermaier S, Schneider T, Bahl MO, Matsubara S, Huesgen PF (2020) Photoprotective Acclimation of the Arabidopsis thaliana Leaf Proteome to Fluctuating Light. Front Genet 11: 1–15

Nikkanen L, Toivola J, Rintamäki E (2016) Crosstalk between chloroplast thioredoxin systems in regulation of photosynthesis. Plant Cell Environ 39: 1691–1705

Nikkanen L, Toivola J, Trotta A, Manuel, Diaz G, Tikkanen M, Aro E-M, Rintamäki E (2018) Regulation of cyclic electron flow by chloroplast NADPH-dependent thioredoxin system. doi: 10.1101/261560

Ojeda V, Pérez-Ruiz JM, González M, Nájera VA, Sahrawy M, Serrato AJ, Geigenberger P, Cejudo FJ (2017) NADPH Thioredoxin Reductase C and Thioredoxins Act Concertedly in Seedling Development. Plant Physiol 174: 1436– 1448

Okegawa Y, Motohashi K (2015) Chloroplastic thioredoxin m functions as a major regulator of Calvin cycle enzymes during photosynthesis in vivo. Plant Journal 84: 900–913

Okegawa Y, Motohashi K (2020) M-type thioredoxins regulate the PGR5/PGRL1- dependent pathway by forming a disulfide-linked complex with PGRL1. Plant Cell 32: 3866–3883

Parker J, Balmant K, Zhu F, Zhu N, Chen S (2015) CysTMTRAQ - An integrative method for unbiased thiol-based redox proteomics. Molecular and Cellular Proteomics 14: 237–242

Peled-Zehavi H, Avital S, Danon A (2010) Methods of redox signaling by plant thioredoxins. Methods in redox signaling

Pérez-Pérez ME, Florencio FJ, Lindahl M (2006) Selecting thioredoxins for disulphide proteomics: target proteomes of three thioredoxins from the cyanobacterium Synechocystis sp. PCC 6803. Proteomics 6 Suppl 1: 186–195

Pérez-Pérez ME, Mauriès A, Maes A, Tourasse NJ, Hamon M, Lemaire SD, Marchand CH (2017) The Deep Thioredoxome in Chlamydomonas reinhardtii: New Insights into Redox Regulation. Mol Plant 10: 1107–1125

Pérez-Ruiz JM, Guinea M, Puerto-Galán L, Cejudo FJ (2014) NADPH Thioredoxin Reductase C Is Involved in Redox Regulation of the Mg-Chelatase I Subunit in Arabidopsis thaliana Chloroplasts. Mol Plant 7: 1252–1255

Pérez-Ruiz JM, Naranjo B, Ojeda V, Guinea M, Cejudo FJ (2017) NTRC-dependent redox balance of 2-Cys peroxiredoxins is needed for optimal function of the photosynthetic apparatus. Proceedings of the National Academy of Sciences 114: 12069–12074

Reichheld J-P, Khafif M, Riondet C, Droux M, Bonnard G, Meyer Y (2007) Inactivation of thioredoxin reductases reveals a complex interplay between thioredoxin and glutathione pathways in Arabidopsis development. Plant Cell 19: 1851–1865

Richter AS, Peter E, Rothbart M, Schlicke H, Toivola J, Rintamäki E, Grimm B (2013) Posttranslational Influence of NADPH-Dependent Thioredoxin Reductase C on Enzymes in Tetrapyrrole Synthesis. Plant Physiol 162: 63–73

Scheibe R (1991) Redox-modulation of chloroplast enzymes: A common principle for individual control. Plant Physiol 96: 1–3

Schjoerring JK, MäcK G, Nielsen KH, Husted S, Suzuki A, Driscoll S, Boldt R, Bauwe H (2006) Antisense reduction of serine hydroxymethyltransferase results in diurnal displacement of NH4+ assimilation in leaves of Solanum tuberosum. Plant Journal 45: 71–82

Schürmann P, Buchanan BB (2008) The ferredoxin/thioredoxin system of oxygenic photosynthesis. Antioxid Redox Signal 10: 1235–74

Selinski J, Scheibe R (2019) Malate valves: old shuttles with new perspectives. Plant Biol 21: 21–30

Serrato AJ, Pérez-Ruiz JM, Spínola MC, Cejudo FJ (2004) A novel NADPH thioredoxin reductase, localized in the chloroplast, which deficiency causes hypersensitivity to abiotic stress in Arabidopsis thaliana. J Biol Chem 279: 43821–43827

Shin JS, Kim SY, So WM, Noh M, Yoo KS, Shin JS (2020) Lon domain-containing protein 1 represses thioredoxin y2 and regulates ROS levels in Arabidopsis chloroplasts. FEBS Lett 594: 986–994

Simionato D, Basso S, Zaffagnini M, Lana T, Marzotto F, Trost P, Morosinotto T (2015) Protein redox regulation in the thylakoid lumen: The importance of disulfide bonds for violaxanthin de-epoxidase. FEBS Lett 589: 919–923

Somerville CR, Ogren WL (1980) Inhibition of photosynthesis in. Nature 286: 257– 259

Stöcker S, Maurer M, Ruppert T, Dick TP (2018) A role for 2-Cys peroxiredoxins in facilitating cytosolic protein thiol oxidation. Nat Chem Biol 14: 148–155

Suzuki A, Knaff DB (2005) Glutamate synthase: Structural, mechanistic and regulatory properties, and role in the amino acid metabolism. Photosynth Res 83: 191–217

Teh JT, Leitz V, Holzer VJC, Neusius D, Marino G, Meitzel T, García-Cerdán JG, Dent RM, Niyogi KK, Geigenberger P, et al (2023) NTRC regulates CP12 to activate Calvin–Benson cycle during cold acclimation. Proceedings of the National Academy of Sciences. doi: 10.1073/pnas.2306338120

Thormählen I, Meitzel T, Groysman J, Öchsner AB, von Roepenack-Lahaye E, Naranjo B, Cejudo FJ, Geigenberger P (2015) Thioredoxin f1 and NADPH-dependent thioredoxin reductase C have overlapping functions in regulating photosynthetic metabolism and plant growth in response to varying light conditions. Plant Physiol 169: pp.01122.2015

Thormählen I, Ruber J, Von Roepenack-Lahaye E, Ehrlich SM, Massot V, Hümmer C, Tezycka J, Issakidis-Bourguet E, Geigenberger P (2013) Inactivation of thioredoxin f1 leads to decreased light activation of ADP-glucose pyrophosphorylase and altered diurnal starch turnover in leaves of Arabidopsis plants. Plant Cell Environ 36: 16–29

Thormählen I, Zupok A, Rescher J, Leger J, Weissenberger S, Groysman J, Orwat A, Chatel-Innocenti G, Issakidis-Bourguet E, Armbruster U, et al (2017) Thioredoxins Play a Crucial Role in Dynamic Acclimation of Photosynthesis in Fluctuating Light. Mol Plant 10: 168–182

Topf U, Suppanz I, Samluk L, Wrobel L, Böser A, Sakowska P, Knapp B, Pietrzyk MK, Chacinska A, Warscheid B (2018) Quantitative proteomics identifies redox switches for global translation modulation by mitochondrially produced reactive oxygen species. Nat Commun. doi: 10.1038/s41467-017-02694-8

Uhrig RG, Echevarría-Zomeño S, Schlapfer P, Grossmann J, Roschitzki B, Koerber N, Fiorani F, Gruissem W (2021) Diurnal dynamics of the Arabidopsis rosette proteome and phosphoproteome. Plant Cell Environ 44: 821–841

Vaseghi M-J, Chibani K, Telman W, Liebthal MF, Gerken M, Schnitzer H, Mueller SM, Dietz K-J (2018) The chloroplast 2-cysteine peroxiredoxin functions as thioredoxin oxidase in redox regulation of chloroplast metabolism. Elife. doi: 10.7554/eLife.38194

Wang P, Liu J, Liu B, Da Q, Feng D, Su J, Zhang Y, Wang J, Wang H bin (2014) Ferredoxin:Thioredoxin reductase is required for proper chloroplast development and is involved in the regulation of plastid gene expression in Arabidopsis thaliana. Mol Plant 7: 1586–1590

Wang P, Liu J, Liu B, Feng D, Da Q, Wang P, Shu S, Su J, Zhang Y, Wang J, et al (2013) Evidence for a role of chloroplastic m-type thioredoxins in the biogenesis of photosystem II in arabidopsis. Plant Physiol 163: 1710–1728

Wittmann D, Geigenberger P, Grimm B (2023) NTRC and TRX-f Coordinately Affect the Levels of Enzymes of Chlorophyll Biosynthesis in a Light-Dependent Manner. Cells 12: 1670

Wong JH, Cai N, Balmer Y, Tanaka CK, Vensel WH, Hurkman WJ, Buchanan BB (2004) Thioredoxin targets of developing wheat seeds identified by complementary proteomic approaches. Phytochemistry 65: 1629–1640

Yamamoto HY, Kamite L (1972) The effects of dithiothreitol on violaxanthin de-epoxidation and absorbance changes in the 500-nm region. Biochimica et Biophysica Acta (BBA) - Bioenergetics 267: 538–543

Yamazaki D, Motohashi K, Kasama T, Hara Y, Hisabori T (2004) Target Proteins of the Cytosolic Thioredoxins in Arabidopsis thaliana. Plant Cell Physiol 45: 18–27

Yano H, Wong JH, Lee YM, Cho MJ, Buchanan BB (2001) A strategy for the identification of proteins targeted by thioredoxin. Proc Natl Acad Sci U S A 98: 4794–4799

Yoshida K, Hara S, Hisabori T (2015) Thioredoxin selectivity for thiol-based redox regulation of target Proteins in Chloroplasts. Journal of Biological Chemistry 290: 14278–14288

Yoshida K, Hisabori T (2016) Two distinct redox cascades cooperatively regulate chloroplast functions and sustain plant viability. Proceedings of the National Academy of Sciences 113: E3967–E3976

Yoshida K, Noguchi K, Motohashi K, Hisabori T (2013) Systematic exploration of thioredoxin target proteins in plant mitochondria. Plant Cell Physiol 54: 875–892

Yoshida K, Yokochi Y, Tanaka K, Hisabori T (2022) The ferredoxin/thioredoxin pathway constitutes an indispensable redox-signaling cascade for light-dependent reduction of chloroplast stromal proteins. Journal of Biological Chemistry 298: 102650

Yu A, Xie Y, Pan X, Zhang H, Cao P, Su X, Chang W, Li M (2020) Photosynthetic Phosphoribulokinase Structures: Enzymatic Mechanisms and the Redox Regulation of the Calvin-Benson-Bassham Cycle. Plant Cell 32: 1556–1573

Zaffagnini M, Fermani S, Marchand CH, Costa A, Sparla F, Rouhier N, Geigenberger P, Lemaire SD, Trost P (2019) Redox Homeostasis in Photosynthetic Organisms: Novel and Established Thiol-Based Molecular Mechanisms. Antioxid Redox Signal 31: 155–210

Zimmer D, Swart C, Graf A, Arrivault S, Tillich M, Proost S, Nikoloski Z, Stitt M, Bock R, Mühlhaus T, et al (2021) Topology of the redox network during induction of photosynthesis as revealed by time-resolved proteomics in tobacco. Sci Adv. doi: 10.1126/sciadv.abi8307

